# The formal demography of kinship. VII. Lifetime kin overlap within and across generations

**DOI:** 10.1101/2025.03.05.641714

**Authors:** Hal Caswell, Charlotte de Vries

## Abstract

**Background:** Interactions among kin have important consequences, including resource transfers, alloparenting, health care, and economic support. Some interactions require that the lives of the interacting relatives overlap. The overlap over a lifetime (lifetime kin overlap, LKO) depends on mortality (longer lives give more opportunity for overlap) and fertility (higher fertility produces more kin with which to overlap). Here we provide a general solution to the problem of calculating lifetime kin overlap.

**Objectives:** To develop a demographic model for the mean and variance of the lifetime overlap of any types of kin over the life of a focal individual.

**Methods:** The matrix kinship model is used to provide the age distribution of kin as an age-specific property of Focal. The mean and variance of lifetime overlap with kin of any type are then calculated using Markov chains with rewards.

**Results:** We obtain the statistics of LKO with numbers of kin, with selected age classes of kin, and with at least one kin, in both prospective and retrospective directions, for each sex separately and with sexes combined. Simultaneous overlap with two or more types of kin (e.g., parents and children) describes sandwich generations. We provide an example comparing Japan under 1947 rates (low survival, high fertility) and 2019 rates (high survival, low fertility).

**Contribution:** It is now possible to compute the mean and variance of the projected LKO with any type of kin, in one-sex or two-sex models based on age or combinations of age and stage.

## 1 Introduction

Interactions among relatives play critical roles in social and family life. Some are within generations and others between generations, and the types of interactions and their effects are diverse. Examples are many; they include resource transfers by bequests (Zagheni and Wagner, 2015; Brennan, James, and Morrill, 1982); support provided by grandparents to children and grandchildren (e.g., Stecklov, 2002; Wachter, 1997; Tu, Freedman, and Wolf, 1993; Himes, 1992), and intergenerational effects on social mobility (Song, 2016; Song and Mare, 2017; Song and Campbell, 2017; Mare and Song, 2015). The presence of relatives can affect infant and child survival (Sear and Mace, 2008) particularly when relatives act as alloparents (Hrdy, 2009). The provision for care across generations imposes costs on those providing the care, particularly in the case of “sandwich” families, in which individuals care for both dependent children and aging parents (DeRigne and Ferrante, 2012; Alburez-Gutierrez, Mason, and Zagheni, 2021). Orphanhood, and more generally bereavement, can be a major disruption of inter-generational interactions during pandemics (e.g., Zagheni, 2010; Snyder et al., 2022). A rich anthropological literature on grandmothers documents their role in caring for grandchildren and the role of such care in the evolution of the human post-menopausal lifespan (e.g., Voland, Chasiotis, and Schiefenhovel, 2005; Hrdy, 2005; Tanskanen and Danielsbacka, 2019). Page and French (2020) have pointed to the importance of kinship structures in determining the balance of kin- and non kin-selection in hunter-gatherer populations. In his PAA presidential address, Mare (2011) emphasized the many possibilities for kin effects among more distantly related individuals, effects encompassing two, three, or more generations.

Some of these interactions can take place only if the participants are alive at the same time; i.e., that their lives overlap. The amount of kin overlap to be experienced over a lifetime depends on the schedules of mortality and fertility, and this has led naturally to attempts to calculate the overlap implied by such a set of demographic rates. We refer to this as ‘Lifetime Kin Overlap,’ abbreviated as LKO.^1^ As we will see, ‘overlap’ can be defined in many ways.

The most sophisticated analysis of LKO to date is that of Song and Mare (2019) who used the kinship model of Goodman, Keyfitz, and Pullum (1974) to calculate the expected lifetime overlap of grandparents with their grandchildren from both prospective and retrospective points of view. Other approaches have been used. Margolis (2016) calculated the time spent with at least one grandchild from prevalence data using the Sullivan method. Microsimulations have been used by Verdery and Margolis (2017) to project older adults without close kin in the United States and by Margolis and Verdery (2019) for a detailed analysis of grandparenthood. Our goal here is to provide a general solution to the lifetime overlap problem. That solution requires two demographic calculations. First, we need to know the nature of the kinship network experienced by an individual of any specified age. For example, a very young person will have no children, but is likely to have parents and even grandparents. An old person is more likely to have children and grandchildren, but unlikely to have living parents and grandparents. The kinship network is a high-dimensional age-specific property of an individual.

Second, we need to integrate this age-specific property over the life on an individual. The individual overlaps, at every age, with a set of kin, up until death. LKO is the lifetime accumulation of this overlap; it reflects the kin available at each age and the probability of living through that age.

The age-specific kinship network is provided by the matrix kinship model. That model has been presented in a series of papers (Caswell, 2019, 2020; Caswell and Song, 2021; Caswell, 2022; Caswell, Verdery, and Margolis, 2023; Caswell, 2024) and is implemented in an R package for those desiring such (Williams et al., 2023). It has been applied to analysis of demographic transitions (Jiang et al., 2023), projections of future kinship networks (Alburez-Gutierrez, Williams, and Caswell, 2023), to prevalences of unemployment (Song and Caswell, 2022) and dementia (Feng, Song, and Caswell, 2024), and even to African elephant social interactions (Croll and Caswell, 2025). In this paper we will use a two-sex version of the model that can, if desired, compute the LKO of female and male individuals with female and male kin (Section 2).

Given the age-specific overlap, the lifetime overlap is calculated using a Markov chain with rewards (MCWR). The MCWR treats the kinship network experienced at each as a ‘reward’ and computes the mean, variance (and higher moments if desired) of the lifetime accumulation of this reward (Caswell, 2011; van Daalen and Caswell, 2017) to give the LKO.

The minimal data needed for these calculations are a mortality schedule and a fertility schedule. If separate schedules are available for men and women, so much the better, but approximations are available if that is not the case. In the example we will explore here, we use separate male and female mortality schedules, which are often available, but only a single female fertility schedule, because male and female fertility are not always available.

Our analyses:

- apply to any type of kin,
- include both prospective and retrospective LKO, in the sense of Song and Mare (2019),
- are easily applied to chosen age groups of kin (e.g., parents older than some age, or children younger than some age),
- accommodate weighted numbers of kin (e.g., kin numbers weighted by prevalence of a disease) and overlap with at least one kin,
- are readily applied to multistate models (e.g., age×parity),
- are flexible in their definition of ‘lifetime,’ including LKO from birth to death (lifetime sensu stricto), remaining LKO starting from any initial age, and LKO up to some specified age (e.g., overlap prior to retirement age),
- provide ‘sandwich’ overlap with two or more types of kin,
- provide not only the mean LKO but also the variance and other statistics,
- make it possible to partition variance in LKO into within- and between-group components.

## 2 The matrix model for the kinship network

The matrix kinship model, which we describe briefly here, projects the expected age distribution of male and female relatives of each of the types of kin alive at each age of a focal individual, referred to as Focal. See Figure 1 for the types of kin. For details on the two-sex version we use, see Caswell (2022).

**Figure 1:**
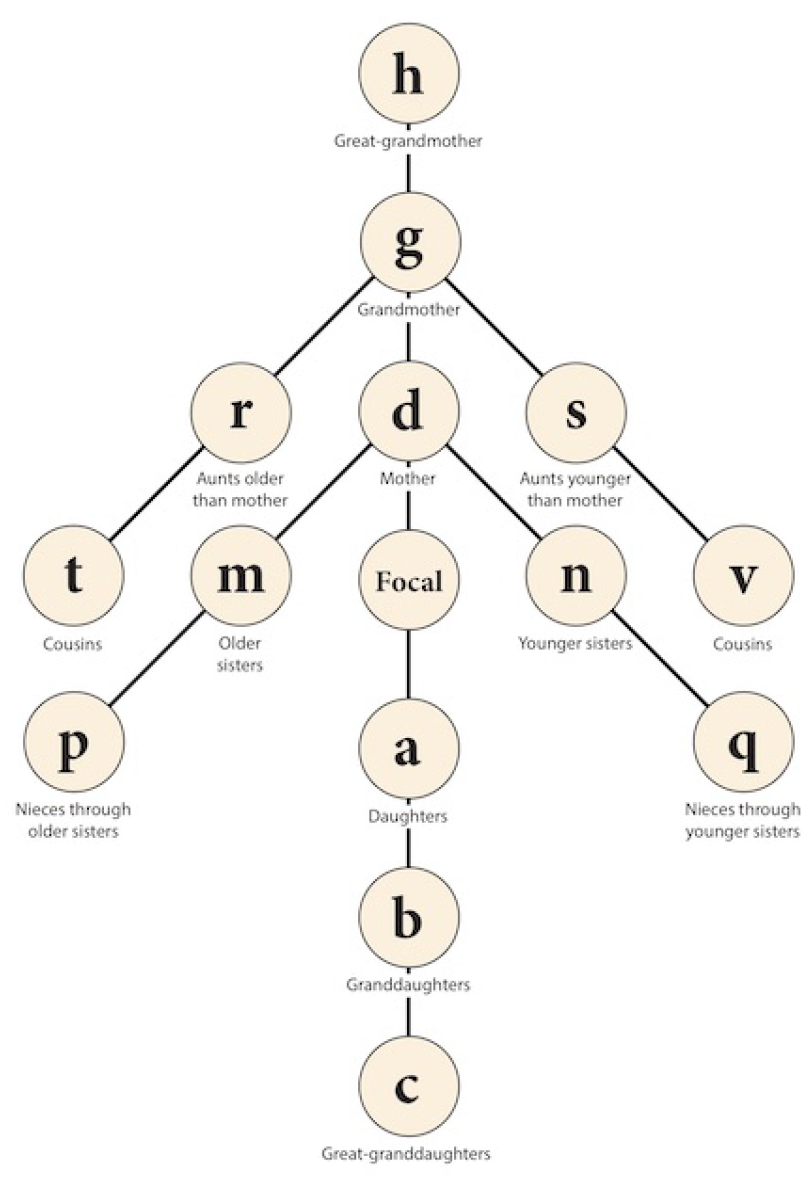
The kinship network showing kin of Focal and the symbols used to identify them. Reproduced from Caswell (2019) under a CC-By license.

### 2.1 Notation

Matrices are denoted by upper case bold characters (e.g., **U**) and vectors by lower case bold characters (e.g., **a**). Vectors are column vectors by default; **x**^T^ is the transpose of **x**. The *i*th unit vector (a vector with a 1 in the *i*th location and zeros elsewhere) is **e**_*i*_. The vector **1** is a vector of ones, and the matrix **I** is the identity matrix. When necessary, subscripts are used to denote the size of a vector or matrix; for example, **I**_*ω*_ is an identity matrix of size *ω* × *ω*. The notation ∥**x**∥ denotes the 1-norm of **x** (i.e., the sum of the absolute values of the entries).

The symbol ◦ denotes the Hadamard, or element-by-element product. On occasion, MATLAB notation will be used to refer to rows and columns; e.g., **F**(*i*, :) and **F**(:, *j*) refer to the *i*th row and *j*th column of the matrix **F**.

Matrices and vectors with a tilde (e.g., Ũ ,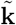) are block-structured in some way, which is specified when they are defined.

### 2.2 Projecting the kinship network: one sex

The kinship network considered here is shown in Figure 1. The network can be extended to longer chains of descendants if desired. In the one-sex model, each type of kin is denoted by a letter, and the bold-faced letter denotes the age distribution vector of that type of kin. For example, **a**(*x*) denotes the age distribution of the daughters of Focal at age *x* of Focal.

The kin of Focal are a population, which is projected by a survival matrix **U** and a fertility matrix **F**. For three age classes,

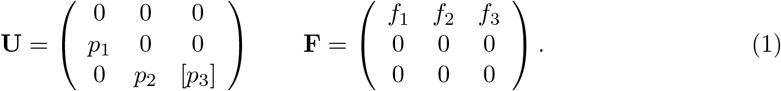

The entry in the lower right corner of **U** is an optional open-ended final age class.

Let **k**(*x*) be the age distribution vector of a generic kin type at age *x* of Focal. The kinship model projects **k**(*x*) by

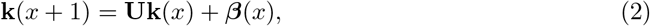

where ***β***(*x*) is a recruitment vector. For some types of kin, no recruitment is possible (e.g., Focal can accumulate no new older sisters). For others, recruitment comes from the fertility of another type of kin (e.g., new granddaughters come from the reproduction of daughters). Thus

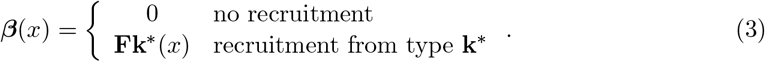

The model is supplemented by an initial condition **k**_0_ that specifies the kin present when Focal is born into the first age class. This is zero for some types of kin (e.g., Focal has no children when she is born), and is calculated for other types from the distribution of ages of mothers at birth. See Caswell (2019) for details.

### 2.3 Projecting the kinship network: two sexes

The one-sex version of the kinship model (Caswell, 2019) describes female kin through female lines of descent. To fully account for both female and male kin (e.g., both grandsons and granddaughters) through all lines of descent (e.g., sons of daughters, sons of sons, daughters of daughters, and daughters of sons) would require both male and female mortality and fertility schedules (Caswell, 2022). In the absence of male and female fertility, we use a model with separate male and female survival, but with female fertility applied to both males and females. This approximation is called Model 2 in Section 5 of Caswell (2022)).^2^

In the two-sex model, the age distribution vector includes both females and males

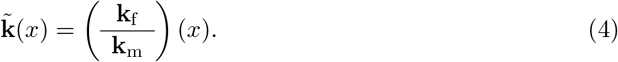

The survival and fertility matrices for Model 2 are block-structured, using female fertility and male and female survival:

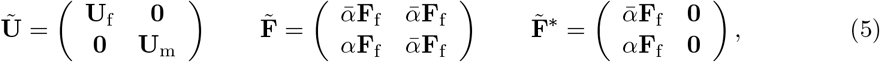

where *α* is the fraction of births that are male and 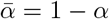. The fertility matrix 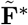 applies to reproduction by direct ancestors (parents, grandparents, and so on). The matrix 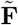 applies to all other kin. See Caswell (2022) for details.

Projecting the kinship model yields the expected age-sex distribution of each type of kin over the life of Focal. This age-sex distribution can be aggregated in various ways (see Section 3.1).

## 3 Markov chain with rewards as a model for overlap

A Markov chain is a stochastic model for the movement of entities among states. In our case, the entities are individuals and the states are a sequence of age classes and one or more absorbing states corresponding to death.^3^ Eventual death is certain. The transitions of an individual are given by a transition matrix^4^

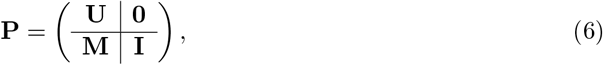

where **U** is the familiar survival matrix (1) and **M** is a matrix containing transitions from each living state to the dead state. The **0** in the upper right corner says that the dead never return to life, and the identity matrix in the lower right corner says that the dead stay dead. The life of an individual is a stochastic sequence of states that continues until eventual death. Although all individuals are identically subject to the probabilities in **P**, some will experience longer, and some shorter lives.

In a Markov chain with rewards (MCWR) the individual collects a ‘reward’ at each step of its life. The reward is a random variable, specified by its moments. Although all individuals collect rewards from the same distributions, some will gather more, and some less, by chance. The individual accumulates the reward over the course of its life until death. The model assumes that the reward, whatever it may be, stops accumulating at death. The variation among individuals in LKO accounts for both sources of stochasticity: the variation of the overlap at each age and the variation in the length of life over which overlap is accumulated.^5^

The age-specific rewards are incorporated into a set of reward matrices. Let *r*_*ij*_ denote the random reward accumulated when an individual makes the transition *j* ↦ *i*. There is a reward matrix for each of the moments of the reward; **R**_1_ contains the first moments, **R**_2_ the second moments, and so on:

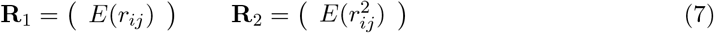

The higher moment matrices are defined similarly. See van Daalen and Caswell (2017) for details on specifying reward moments.

The lifetime accumulation of rewards is given in a set of vectors, 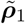 for the first moments and 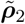 for the second moments. The *i*th entry of 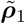 is the mean remaining lifetime reward for an individual in age class *i*; similarly for the second moments in 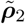. The formulae for these moment vectors are given by van Daalen and Caswell (2017, Theorem 1). In our case they are block-structured for males and females:

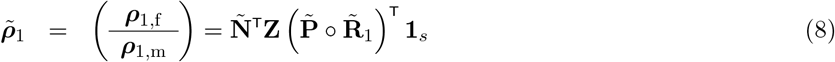

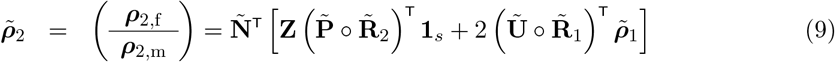

where Ñ is the fundamental matrix,

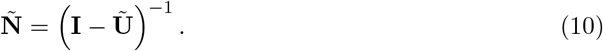

The matrix **Z** is a matrix of zeros and ones that slices off the rows and columns of 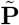 and 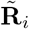 corresponding to the absorbing states.

The first two moments suffice to calculate the mean, variance, standard deviation, and coefficient of variation of LKO:

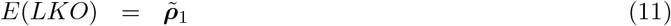

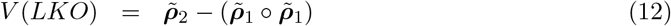

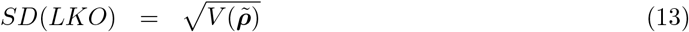

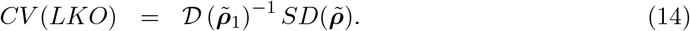

The means, variances, and so on are taken element-wise. These quantities are vectors whose entries give the statistics of remaining LKO.

### 3.1 Measures of age-specific overlap

When the question is posed about overlap with some type of kin, one must specify the measure of kin abundance that is of interest. Do we want to know overlap with all grandchildren? Or with young grandchildren for whom you might babysit? Or with teen grandchildren who you might help with education? Whatever the choice, we denote this measure as *ξ*. Some possibly interesting choices include

- The age distributions of female and male kin are given by the blocks **k**_f_(*x*) and **k**_m_(*x*) in 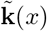 . To calculate LKO with both sexes combined, we define the age-specific overlap as

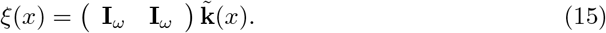
- Overlap with female and of male kin separately are given by

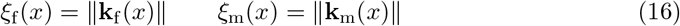

and the total number of kin, female and male combined, is

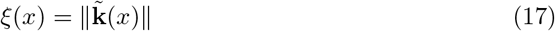
- We might be interested in overlap with the number of kin in selected age ranges (e.g., school age children); this is given by using as the measure of abundance

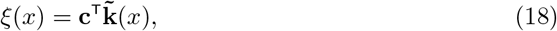

where **c** is a 0-1 vector that selects the age classes and sexes of interest.
- We might be interested in overlap with kin weighted by some measure of importance. If **w** is a vector of age-specific weights, then

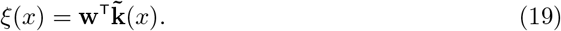

Weights might be measures of prevalence of, for example, employment, disability, health conditions, or as in Song and Mare (2019) grandparents weighted by educational attainment. In an evolutionary context, kin might be weighted by their degree of relatedness to the focal individual; overlap with more closely related kin permits more intense selection on traits related to kin interactions.

### 3.2 Reward matrices for kin overlap

The matrix containing the first moments of age-specific overlap is

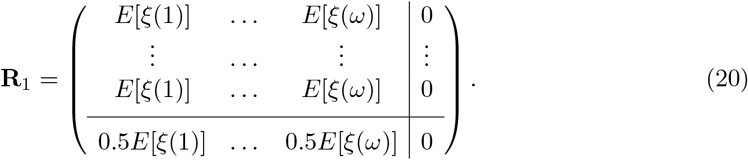

The entries in the last row are the rewards collected if Focal dies during that year, assuming that on average, she lives half the time step).^6^

The matrices for higher moments depend on the frequency distribution of kin numbers. These results are provided by the stochastic kinship model (Caswell, 2024). Here, we will use the fact that the numbers of kin are well fit by a Poisson distribution except for direct ancestors (parents, grandparents, and great-grandparents), which are well described by a binomial distribution. Both of these distributions provide variances for the kin numbers at every age; the variance of the age-specific overlaps in Section 3.1 can be calculated from these. The second moment reward matrix is then

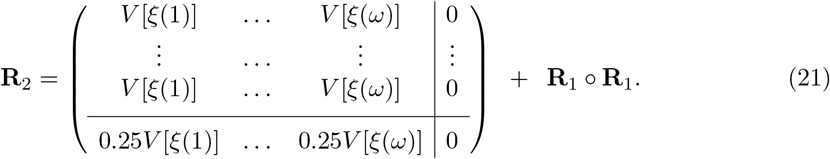

### 3.3 LKO is measured in units of person-years

Place yourself, for a moment, in the shoes of an individual progressing through their life. At each age your life overlaps with some number of, say, cousins. As each year goes by, you accumulate more overlap, and add that years overlap to the overlap already accumulated. LKO thus has units of person-years. This implies that spending one year with 10 cousins is equivalent, as far as you are concerned, to spending 10 years with one cousin.

An alternative would be to ask, in each year, “do I have any cousins?” and to accumulate overlap with at least one cousin (see Section 4.2). This version of LKO is measured in years.^7^ Song and Mare (2019) referred to these two choices as “overlap with all kin” and “overlap with any kin”.

## 4 Kin overlap during a demographic transition in Japan

As an example of the LKO calculations and their results, we present an analysis of kin overlap for the population of Japan (as in Caswell 2019 and Caswell 2024). Japan underwent a dramatic demographic transition between 1947 and 2019. Under 1947 rates, period life expectancy in Japan was low (53.7y) and fertility was high (period TFR of 4.6). By 2019, Japan had one of the highest life expectancies in the world (87.4y) and one of the lowest fertilities (TFR of 1.3). This dramatic transition translates into important differences in the kinship network (Caswell, 2019, 2024).

As an example, expected LKO with grandchildren and grandparents is shown in Figure 2. Lifetime overlap at birth with grandchildren is higher under 1947 rates than under 2019 rates (about 125 person-years compared to 50 person-years). LKO with grandparents is about 45 person-years under 1947 rates and about 100 person-years under 2019 rates.

**Figure 2:**
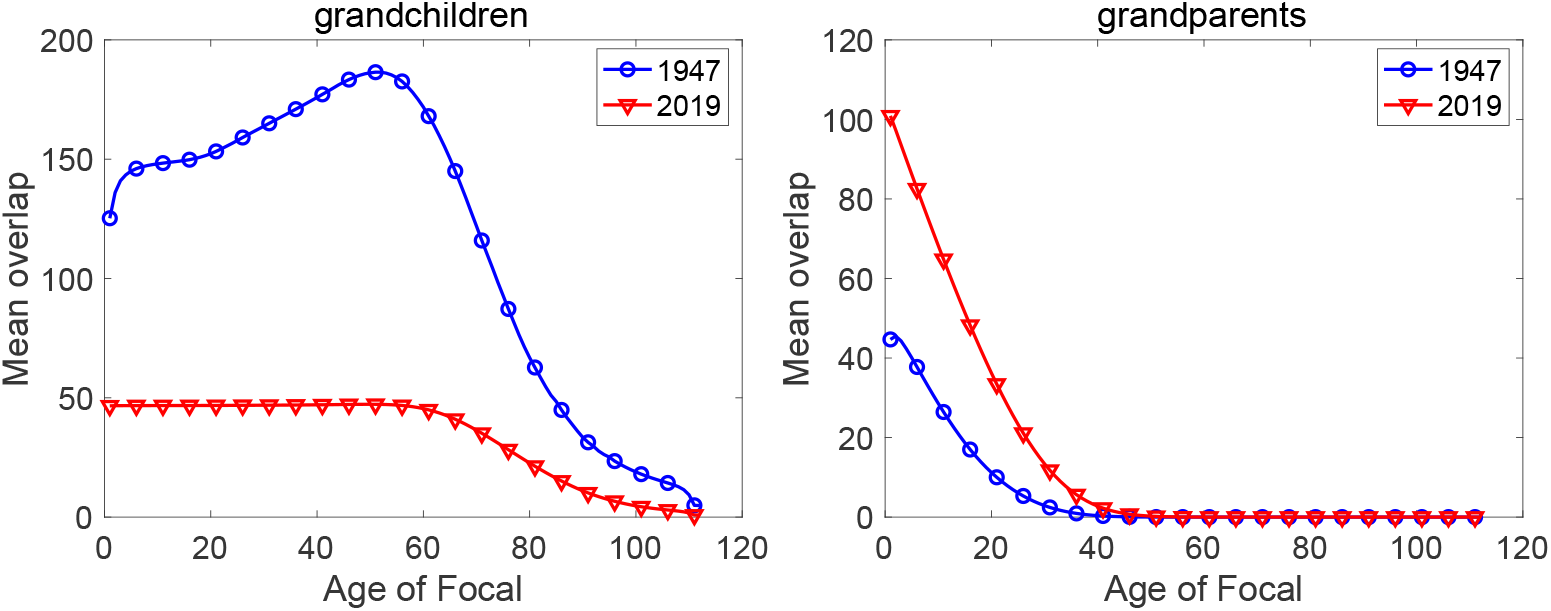
Mean remaining lifetime overlap with grandchildren and grandparents, in person-years, as a function of the age of Focal. Japan data for 1947 and 2019; male and female kin combined. Note different scales on the *y*-axes.

**Collected figures for all kin.** For convenience, figures showing the LKO results for all types of kin are collected in Appendix A. For ready reference, the expected kin numbers for each type of kin, under the 1947 and 2019 rates, are given in Appendix B.

### 4.1 Lifetime, remaining lifetime, and partial lifetime overlap

The vectors 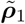 and 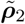 defined in Equations (8) and (9) give the statistics of remaining lifetime overlap, starting from every age. Partial LKO, up to some specified age *x*_max_ rather than to death, is calculated by terminating the overlap calculation at *x*_max_. This can be done in two ways. One is to artificially ‘kill’ Focal at age *x*_max_ by setting the survival probability used in the MCWR calculation (the subdiagonal entries in **U** in Equation (6)) to 0 for ages *x > x*_max_. Equivalently, one could set the reward matrices to zero for ages greater than *x*_max_,

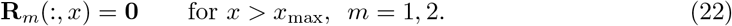

This would allow Focal to live beyond age *x*_max_, but would prevent the accumulation of further rewards after that age.

Mean LKO results for all types of kin are collected in Figure A-1. As with grandparents in Figure 2, LKO with parents and great-grandparents is higher under 2019 rates than under 1947 rates, reflecting the higher survival in 2019. LKO with all other types of kin is higher under 1947 rates, reflecting the higher fertility in 1947.

### 4.2 Overlap with at least one kin

Instead of focusing on age-specific overlap with the number of kin, consider the question of overlap with at least one relative (e.g., at least one child, or at least one sibling). Song and Mare (2019) refer to this as overlap with ‘any kin’. This calculation treats the overlap as identical regardless of the number of children or siblings involved.

The condition of having at least one relative is a prevalence measure, formally similar to the condition of having a disability or a disease. Our task is to calculate the age-specific prevalence of having at least one relative, given the mean and variance in the number of kin provided by the kinship model. This requires information on the probability distribution of kin numbers at each age. Even if the expected number of kin is less than one, there is still some probability that at least one kin is present. Similarly, even if the expected number is large, there is still some probability to have none.

Given the prevalence of at least one kin, we then compute the reward matrices **R**_1_ and **R**_2_. See Caswell and Zarulli (2018) and Zarulli and Caswell (2024) for examples of computing reward matrices for the prevalences of health conditions.

If the number of kin follows a Poisson distribution, then the prevalence of having at least one relative at age *j* is

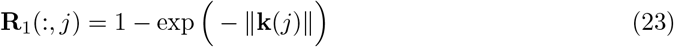

The Poisson distribution is appropriate for all categories of kin except for direct ancestors (parents **d**(*x*), grandparents **g**(*x*), and great-grandparents **h**(*x*) in the current model). For them, the appropriate distribution of kin numbers is binomial, with sample sizes *N* = 2, 4, 8, respectively (see Caswell, 2024). The prevalence of having at least one relative in the binomial case is

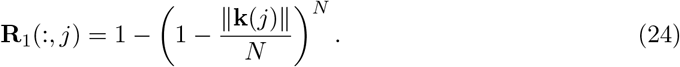

Because a relative is either present or not present, the presence of kin has a Bernoulli (0-1) distribution^8^ and the second moment reward matrix is the same as the first:

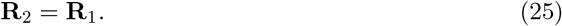

Lifetime overlap with at least one kin calculated from these matrices has units of years.

Figure 3 shows the mean LKO with at least one grandchild and at least one grandparent. Unlike LKO with numbers of grandparents and grandchildren in Figure 2 , LKO with at least one relative is higher under 2019 rates than under 1947 rates. LKO with numbers of grandchildren declines from ∼ 65 person-years under 1947 rates to ∼ 23 person-years under 2019 rates. Overlap with at least one grandchild increases from ∼ 17 years in 1947 to ∼ 25 years in 2019. Similarly, LKO with numbers of grandparents increases from ∼ 20 − 25 person-years in 1947 to ∼ 45 − 55 person-years in 2019. LKO with at least one grandparent increases from 22 years to 38 years over the same interval.

**Figure 3:**
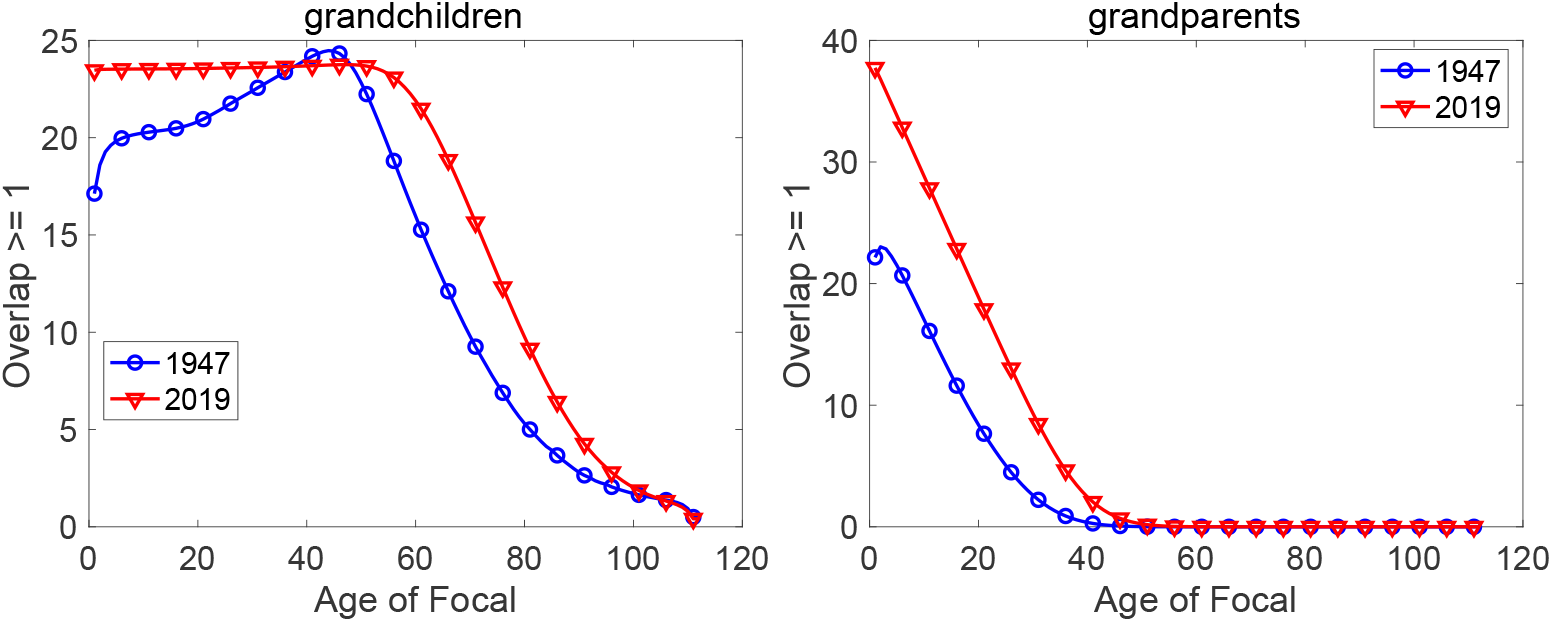
Mean LKO with at least one grandchild (left) and at least one grandparent (right), under Japanese rates for 1947 and 2019, male and female kin combined. Compare with Figure 2, which shows LKO with numbers of kin. Note different y-axis scales.

### 4.3 Overlap with female and male kin

The extent of overlap with female and male kin may differ because of fertility and mortality differences between the sexes. Instead of combining the sexes as in Figure A-1, in Figure A-2 we show results for each sex (i.e., sons and daughters, mothers and fathers, and so on).

The differences in overlap between the sexes are small. They are nearly invisible for children, grandchildren, and great-grandchildren. Even under 1947 rates these kin are unlikely to be old enough for sex differences in longevity to have an impact. The biggest differences between LKO with female and male kin are for parents, grandparents, and great-grandparents. Focal will experience more person-years of LKO with mother than father, with grandmothers than grandfathers, and great-grandmothers than great-grandfathers.

The model in Equations (4) and (5) contains sex differences only in mortality. A model that also included female and male fertility would probably not change these LKO patterns because while births may differ between the sexes, the sex ratio of offspring will not.

### 4.4 Variation and prediction intervals for LKO

The MCWR provide the second moments, and thus the variances and standard deviations of LKO, in Equations (9), (12), and (13). These measure the variation among individuals in their experience of LKO. Figure A-4 shows the standard deviation as a function of the age of Focal for all kin types. These standard deviations are almost always much higher under 1947 than under 2019 rates. Thus, under 1947 rates not only do individuals have higher expected overlap with their families, but there is more variation among individuals in this family experience.

The variance in LKO implies a prediction interval surrounding the expected value.^9^ The interval depends on the probability distribution of lifetime overlap. Kin overlap is highly overdispersed relative to a Poisson distribution, so the gamma distribution, fit by the mean and variance, is an attractive choice. The gamma has support on the nonnegative real line, and includes the exponential, Weibull, *χ*^2^, and Erlang distributions as special cases. The method of moments provides estimates of the shape parameter *a* and the scale parameter *b* of the Gamma distribution of a variable *ξ* as

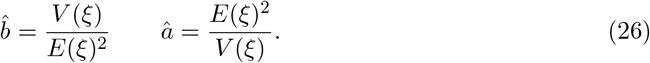

The desired prediction intervals are then obtained from the inverse of the Gamma distribution.

The 90% prediction intervals for LKO with grandchildren and grandparents are shown in Figure 4. The intervals for grandchildren are wide: from about 10 to about 300 person-years under 1947 rates. There is much less variation around the mean for grandparents. For completeness, the prediction intervals for all kin are given in Figure A-5.

**Figure 4:**
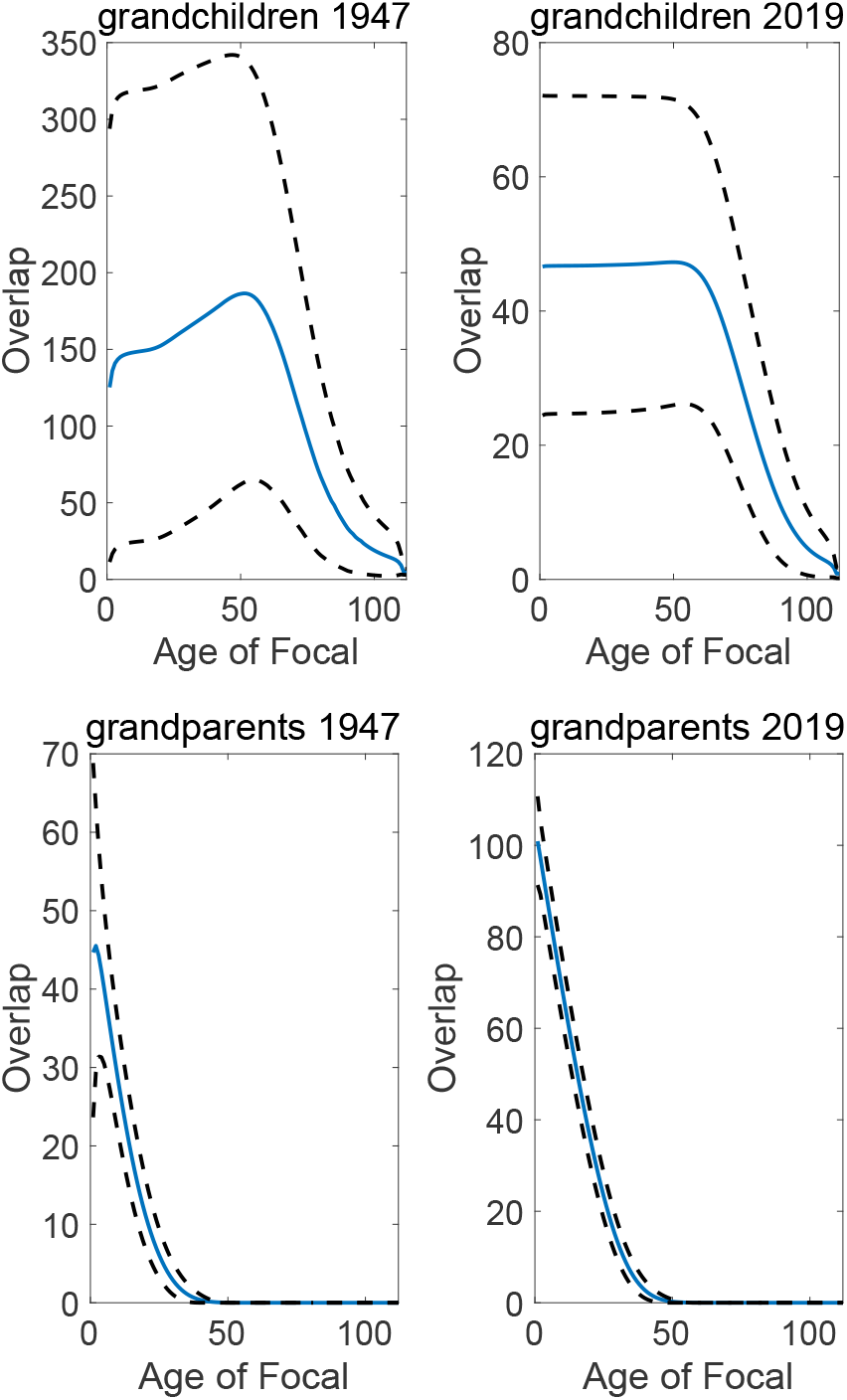
The mean and 90% prediction intervals for LKO with grandparents and grandchildren under Japan rates in 1947 and 2019. Note different *y*-axis scales.

**Figure 5:**
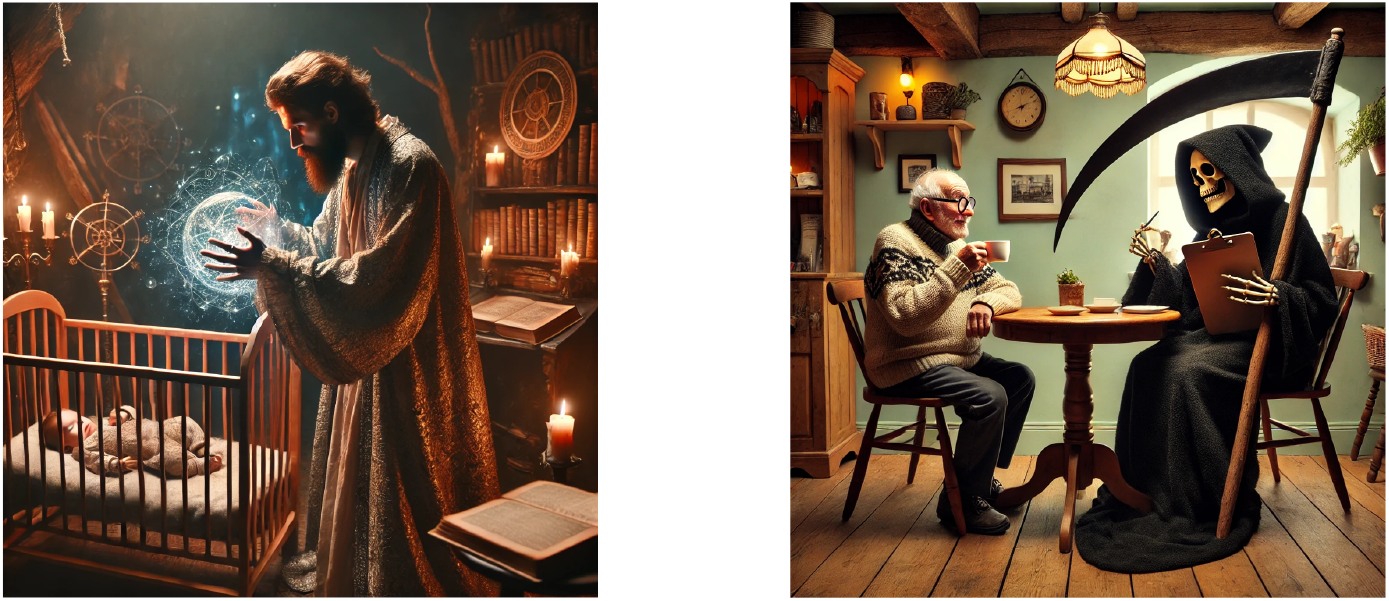
Prospective and retrospective overlap. Images created using ChatGPT.

### 4.5 Prospective and retrospective overlap

Song and Mare (2019) made an important distinction between prospective and retrospective overlap with kin. Consider overlap with grandparents. Prospective LKO projects this overlap forward from Focal’s birth (or some other age) until death, as if with a crystal ball informed by mortality and fertility rates. The calculation must account for both the kin with whom she will overlap at each age and her potential lifetime to experience that overlap. The results we have shown so far are all prospective overlap measures.

Retrospective LKO can be thought of as the response of an individual (let us call her Respondent) at a given age *a* who is confronted with a survey question asking how much overlap she has had with grandparents up to age *a*. The calculation must be conditional on Respondent’s survival to age *a*, so that she is available to respond to the survey question, and must exclude any overlap after age *a*, because Respondent has no knowledge of that.

The calculation of retrospective overlap is straightforward. To condition on Respondent’s survival to age *a*, we modify the **U** matrix in Equations (6), (8), and (9) to contain ones on the subdiagonal and zeros elsewhere.

Because overlap after age *a* has no impact on Respondent’s retrospective overlap up to age *a*, we might as well calculate as if Respondent dies, or stops collecting rewards, immediately after answering the retrospective survey question at age *a*. An easy way to do this is to set the columns of the reward matrices to 0 for all columns greater than *a*

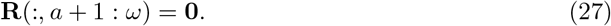

The retrospective LKO of Respondent with her grandchildren and grandparents is shown in Figure 6. Retrospective LKO is by necessity a non-decreasing function of Respondent’s age. The retrospective LKO with grandchildren is zero until Respondent is about age 50, after which it increases with Respondent’s age. The increase is more rapid, and reaches much higher levels, under 1947 rates than under 2019 rates.

**Figure 6:**
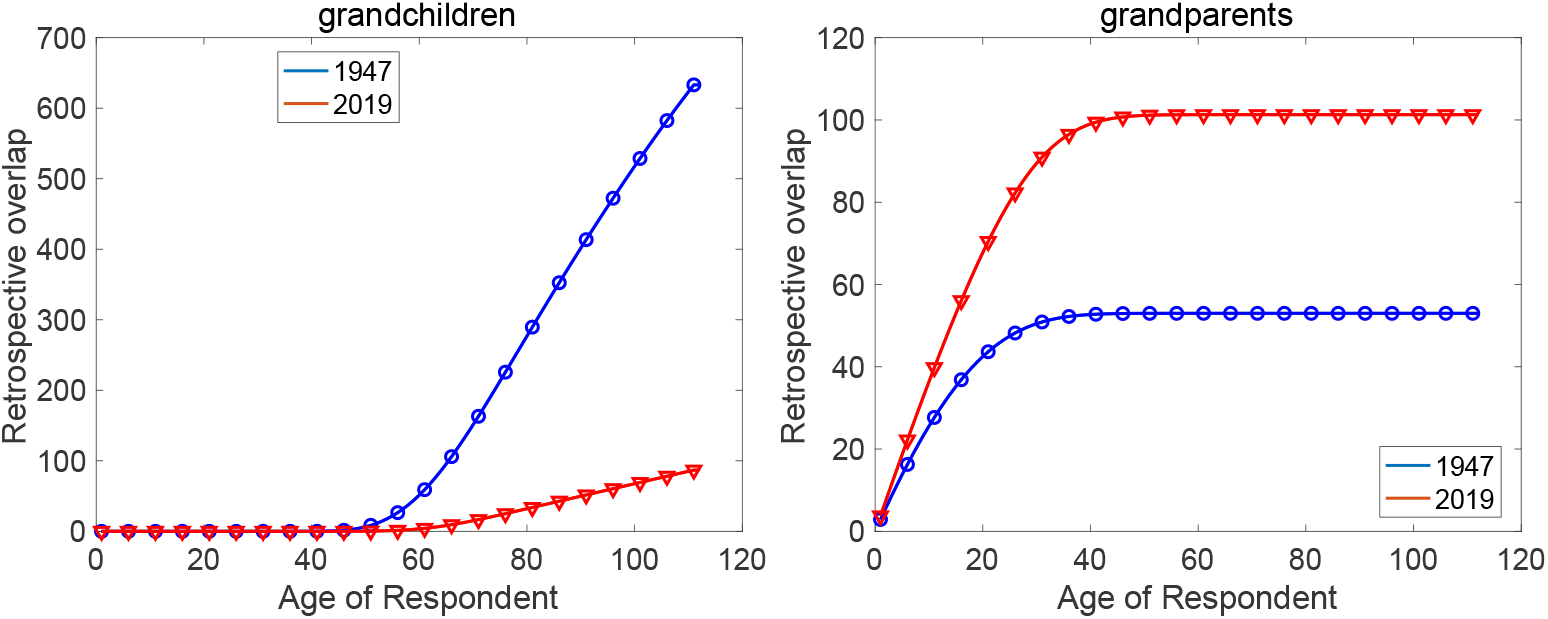
Mean retrospective LKO with grandchildren and grandparents, in person-years, as a function of the age of Respondent. Japan data for 1947 and 2019, both sexes combined. Note different scales on the *y*-axes.

Retrospective LKO with grandparents increases to an asymptote at about age 30 under 1947 rates and about age 40 under 2019 rates. After those ages, Respondent’s grandparents are likely not alive and so Respondent’s LKO with them cannot increase further. The overlap is higher under 2019 rates than under 1947 rates.

Retrospective LKO results for all kin are shown in Figure A-6.

### 4.6 Simultaneous overlap: The case of sandwiched kin

Focal experiences overlaps with many types of kin, and sometimes those simultaneous overlaps create problems or opportunities. The term ‘sandwich generation’ has been introduced to describe an individual sandwiched between simultaneous overlap with, and associated care burdens for, young children and old parents (e.g., Lei, Leggett, and Maust, 2023; DeRigne and Ferrante, 2012; Alburez-Gutierrez, Mason, and Zagheni, 2021).

The sandwich concept can be generalized to other types of kin and need not always have negative connotations of burdens of care. Being sandwiched between young children and parents who are loving grandparents to those children can be a benefit rather than a burden to Focal (e.g., Eibich and Zai, 2024). Similarly, simultaneous overlap of Focal with children and siblings might measure the opportunity for alloparental care of Focal’s children by their aunts or uncles (Nitsch, Faurie, and Lummaa, 2014). Kinlessness is a particular null case of overlap where Focal’s simultaneous overlap with some specified type(s) of kin is zero (e.g., Verdery and Margolis, 2017; Margolis and Verdery, 2017; Margolis et al., 2022).

To be sandwiched at age *x* is to overlap simultaneously, at age *x*, with two types of kin. Consider two types (parents and children, for example) and define weighted kin numbers *ξ*_1_(*x*) and *ξ*_2_(*x*) at each age

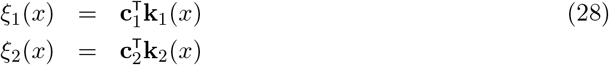

where **c**_1_ and **c**_2_ are vectors of weights, as in Section 3.1.

An appropriate measure of the simultaneous overlap of Focal with two types, at age *x*, is given by the geometric mean^10^ of the two abundances

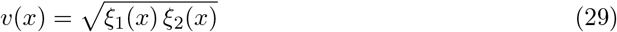

The first moment reward matrix for ‘sandwichedness’ is then

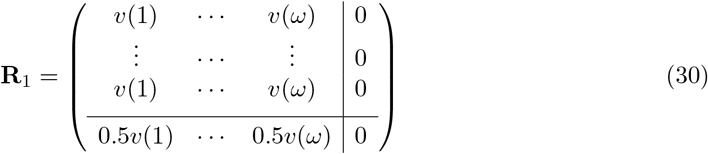

As a measure of simultaneous overlap, the product of the numbers of two types of kin has the desirable property of being zero in the absence of either type of kin, and of being an increasing function of either type of kin for a fixed value of the other. The square root in the geometric mean standardizes the units of *v*(*x*) to individuals rather than individuals squared.

The entries of the vector 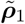 from equation (8) are the mean LKO, in units of sandwiched-person-years, over the lifetime of Focal. The second moment matrix **R**_2_ would require the variance of the geometric mean of kin numbers at each age. This variance could be found using a series expansion, but will not be explored here.

#### 4.6.1 The layers of the sandwich

A variety of definitions of the young and old parts of the sandwich have been proposed. Some analyses define young dependent children as those less than 18 years of age, and old dependent adults as older than 65 years. A particularly interesting definition by Alburez-Gutierrez, Mason, and Zagheni (2021) defines the young as less than or equal to 15, and defines the old as those within 5 years of death, using the prospective longevity approach of Sanderson and Scherbov (2019). We do not explore that definition here, but it will be interesting to incorporate it into the MCWR framework.

It is possible to define sandwichedness in terms of any set of kin. Alburez-Gutierrez, Mason, and Zagheni (2021) examine parent-child sandwiches and parent-grandchild ‘grandsandwiches’. It is possible to define multilayer sandwiches with more than two types of kin, for example simultaneous overlap with parents, siblings, children, and nieces-nephews. This would capture a three-generation family configuration that may be of interest in family care. The geometric mean measure of overlap in such a case would be

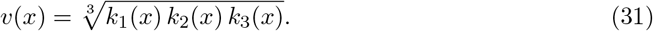

Hünteler (2022) and Hünteler, Nutz, and Wörn (2024) have defined and explored a range of generational family structures, ranging from childlessness through one-, two-, three- and four-generation families. The patterns of LKO in such multi-layer sandwiches would be interesting to explore.

For the case of Japan, Figure 7 shows the expected LKO, in units of person-years, for two definitions of the sandwich under 1947 and 2019 rates. A set of ‘mild’ conditions defines Focal as sandwiched when she overlaps simultaneously with children younger than 18 and parents older than 65. A more strict definition is for simultaneous overlap with children younger than 5 and parents older than 75. The age patterns of LKO with these sandwiches are very similar, but of course the amount of sandwichedness is greater when the definitions are relaxed to include a wider age range of children and parents.

**Figure 7:**
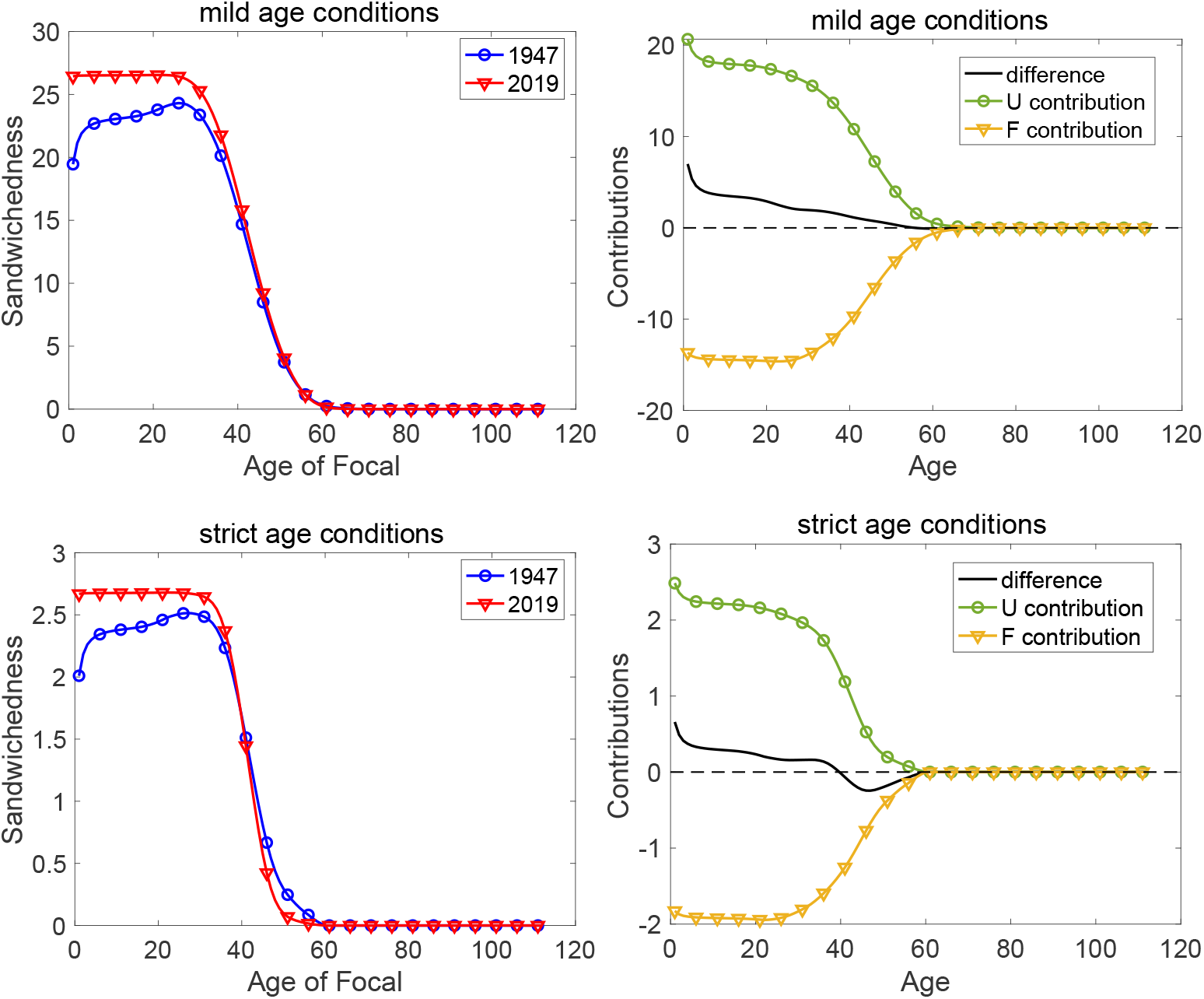
Left: Mean LKO as a sandwich between parents and children under milder (parents older than 65, children younger than 18) and stricter (parents older than 75, children younger than 5) age conditions. Right: Decomposition of the difference between LKO under 2019 and 1947 rates into contributions from the survival and fertility differences.

## 5 Decomposing differences in overlap: Contributions of mortality and fertility

The difference in LKO between two populations (or two time periods) is influenced by both fertility and survival. When both fertility and survival differ, as in the comparison of Japan under 1947 and 2019m rates, the difference in LKO can be decomposed into contributions of mortality and fertility using the classic Kitagawa-Keyfitz decomposition method (Kitagawa, 1955; Keyfitz, 1968)

It is often supposed that sandwiched families will become more common under recent conditions of higher survival and longer life expectancy. However, that has not happened in the case of Japan under 1947 and 2019 rates, despite the dramatic difference in those rates. Under either mild or strict age conditions, the mean sandwiched LKO is quite similar between the 1947 and 2019 rates.

To decompose this difference, consider two populations, each with its own survival and fertility matrices, and let ***ρ***_1_ be the vector of mean LKO sandwichedness calculated using the reward matrix (30). In our case the two populations are Japan in 1947 and in 2019. The difference in overlap is

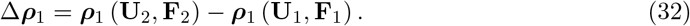

We partition this difference into contributions from the difference in **U** and the difference in **F**. These contributions are

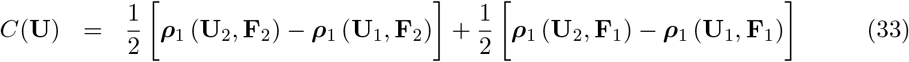

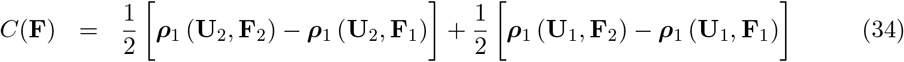

The first term on the right-hand side of *C*(**U**) is the difference in LKO that would be produced by the difference in survival matrices **U**_2_ − **U**_1_ under the fertility schedule **F**_2_. The second term is the same, but under the fertility schedule **F**_1_. The contribution of differences in **U** is the average of these two. The calculation is the same for the effects of the differences in **F**.

If ***ρ***_1_ is a vector (e.g., a vector of remaining lifetime overlap as a function of age), the contributions *C*(**U**) and *C*(**F**) will also be vectors. The decomposition is exact.

The decomposition result applied to Japanese rates in 1947 and 2019 is shown in the right hand panels of (Figure 7). The increased survival in 2019 would, on its own, have created a large increase in sandwiched LKO (+25 person-years with the mild age definition, +3 person-years with the more strict age definition). The reduced fertility in 2019 would, on its own, have created an almost equal reduction in sandwiched LKO. The contributions of survival and fertility differences nearly cancel each other out, leaving little difference in sandwiched LKO.

Is there something special about the comparison of these two particular years, in this particular country, that leads to such a close balance between increases and decreases? We do not know.

## 6 Discussion

Humans are a social species. Our interactions with immediate family, ancestors, descendants, and other relatives affect many aspects of our lives and family dynamics. Many of the interactions require shared lives; they take place only if the lives of both parties overlap. This calls for a way to calculate the patterns of lifetime overlap (LKO) implied by a set of demographic rates. Such calculations have previously been approached by formal models (e.g., Song and Mare, 2019) and by microsimulations (e.g., Margolis and Verdery, 2019; Alburez-Gutierrez, Mason, and Zagheni, 2021). Valuable as those analyses are, they have been limited to a few types of kin and indices of overlap, and have been provided only expected values of kin overlap. A more general and flexible framework, that can incorporate a wider range of demographic outcomes and provide variances as well as means, has been lacking.

### 6.1 Overview

We have approached the problem by combining two analyses: (1) the kinship network surrounding Focal at each age and (2) the life course of Focal as she experiences that network over a lifetime. The first of these is provided by the matrix kinship model, the second by a Markov chain with rewards. The rewards accumulated at each age (or more generally each transition) are given by (some function of) the age-specific kinship network.

The rewards are incorporated in matrices defined in Equations (20) and (21) for kin abundance, by (23) and (24) for presence of at least one kin, and by (30) for sandwiched kin. The MCWR calculations are given by Equations (8) and (9). Higher moments are available if desired.

Our comparison of Japan under 1947 and 2019 rates highlights changes in LKO produced by the dramatic demographic transition that occurred over that period. The rates of 1947 produced much greater LKO with all types of kin except parents, grandparents, and great-grandparents, compared to 2019 rates. In contrast, the rates of 2019 produced greater LKO with at least one relative. The variance in LKO among individuals is much greater under 1947 rates than 2019 rates. The reduction in fertility and increase in survival between 1947 and 2019 work in opposite directions. In the case of sandwich overlap with parents and children, the contribution of these two changes almost cancel each other out.

### 6.2 Evolutionary demography

In evolutionary and anthropological contexts, overlap with kin who have different degrees of genetic relatedness influences the outcome of kin selection (Tanskanen and Danielsbacka, 2019). For example, alloparenting of kin in humans allowed us to have shorter birth intervals when compared to other primates with altricial offspring (Hrdy, 2009). In turn, Hrdy (2009) argues that this complicated and social form of childrearing led to the evolution of our unique prosocial tendencies.

The evolutionary importance of overlap with kin is not limited to humans, but applies more broadly to other social species. For example, Ellis et al. (2024) argued the evolution of menopause in toothed whales (one of the few mammalian groups that share this property with humans) is related to their kinship networks. The toothed whales are projected to have more kin than other groups, increasing the opportunity for post-reproductive females to benefit their relatives and thus increase inclusive fitness. Refining those analyses to consider kin overlap as well as abundance might provide additional insight.

The mortality and fertility of elephants are also influenced by the overlap with relatives (e.g., Lahdenperä et al., 2012; Lahdenperä, Mar, and Lummaa, 2016a,b; Berger et al., 2021; Croll and Caswell, 2025). Analyzing LKO and its response to poaching pressure, as was done for population growth by Croll and Caswell (2025), would be an exciting addition.

The LKO calculation also provides a potential starting point for empirical estimates of the lifetime accumulated benefits due to helping; that is, estimates of one side of Hamilton’s equation (Hamilton, 1964). Consider sibling helping in elephants as an example. Lynch et al. (2019) found that living near a younger sister significantly increased the likelihood of annual reproduction among female elephants, and this effect was strongest when living near a sister 0–5 years younger. The analysis presented here might be used to calculate the lifetime accumulated additional nephews and nieces born due to a focal female overlapping with sisters. Incorporating reproductive value (which describes the relative importance of a stage from the perspective of natural selection; see Rodrigues and Gardner 2022) would allow an estimate of the benefit of sister helping, i.e. the benefit side of Hamilton’s equation. Obtaining empirical estimates of Hamiltons equation has been challenging (see Bourke 2014 and van Veelen et al. 2017 for reviews), so this would be an exciting development.

### 6.3 Extensions

Several issues are not yet addressed in the method we present here. Although male and female rates are included, spouses, blended families, and kin by marriage are not. Including these aspects of kinship networks is an open research problem. However, both Focal and Focal’s spouse are subject to the same demographic rates, so the patterns found here are expected to be mirrored in a model including affinal kin.

Our analysis uses time-invariant period rates, and thus apply to synthetic cohorts following those rates over a lifetime. Extension to time-varying rates is an open question. We have used age-classified demographic rates. Multistate or stage-classified models could be analyzed as was done for age×parity by Caswell (2020), provided that all offspring are born into the same stage. Analysis of cases in which offspring are born into different stages (e.g., spatial models) require further extensions. To date, analyses of such cases require the approach introduced by Coste et al. (2021) and Butterick et al. (2024). Extending the matrix kinship model to cover these cases is an open research problem; see Butterick et al. (2025) for some promising developments.

A dimension of overlap yet to be addressed is spatial proximity, especially coresidence. Incorporating the prevalence of coresidence into a weighted kin abundance measure, as in Equation 19, would be a first step towards examining LKO with age-weighted coresident kin.

The MCWR analysis yields the moments, as many as desired, of the distribution of LKO (van Daalen and Caswell, 2017). We have focused on the mean and variance of LKO here; the skewness of LKO could be calculated from the third moments of LKO. An alternative approach to lifetime calculations, in the context of lifetime reproductive success, was introduced by Tuljapurkar et al. (2020). That approach provides the complete distribution; it may be possible to extend this approach from lifetime reproductive output to LKO.

We have indicated some of the potential applications of this analysis, and shown some examples of how LKO changes over a demographic transition. More patterns await exploration.

## 7 Acknowledgements

This research was supported by the European Research Council under the European Union’s Horizon 2020 research and innovation program, ERC Advanced Grant 788195 (FORMKIN) to HC. LdV was also supported by Swiss National Science Foundation grant P500PB 211003 and by NWO grant VI.Veni.222.400. We thank Xi Song for helpful comments and particularly influential work on kin overlap.

## Appendix A Collected figures: LKO results for all kin

### A.1 Mean prospective lifetime overlap with kin

**Figure A-1:**
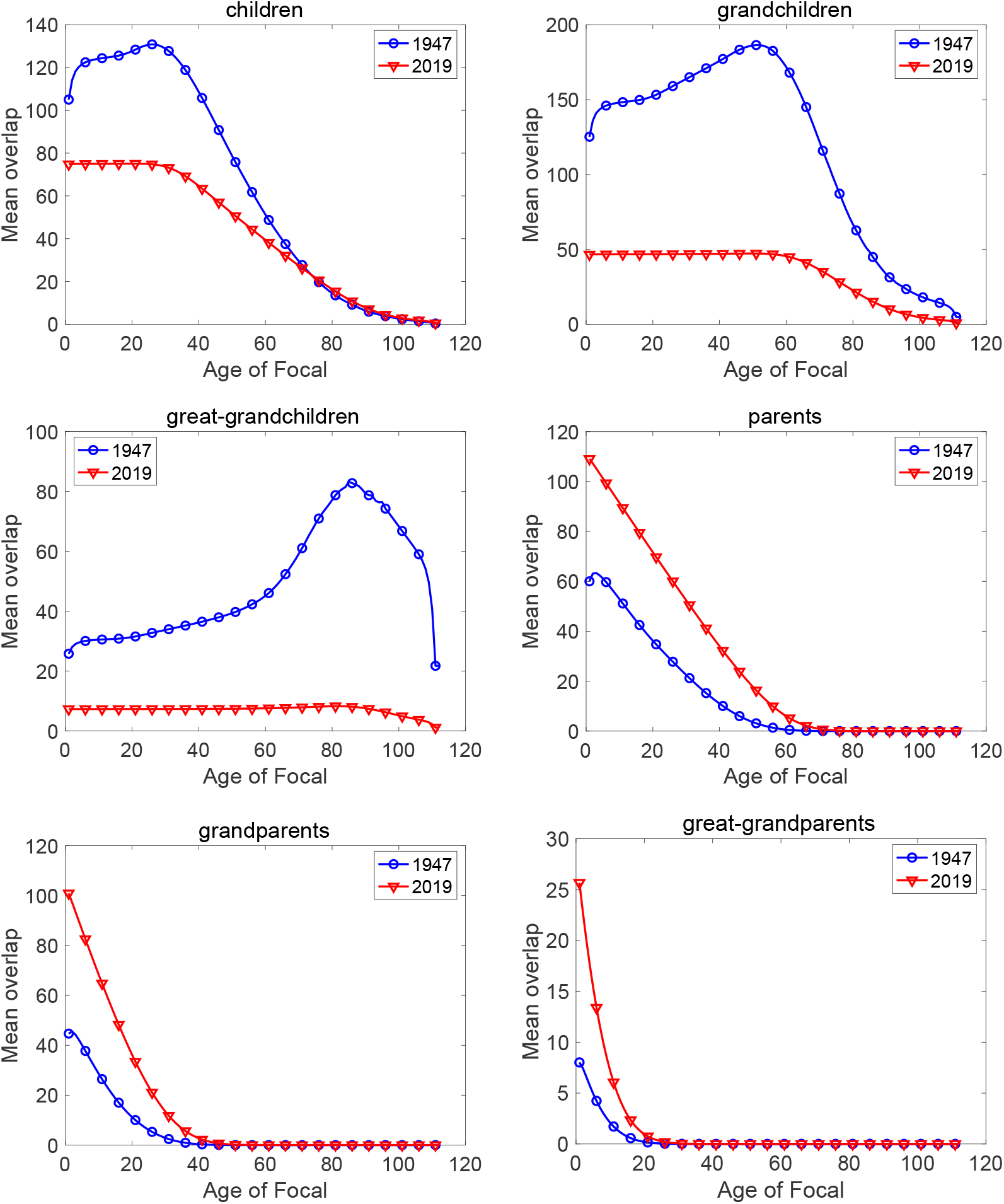
(**Part 1**) Mean remaining LKO with each type of kin, both sexes combined, as a function of the age of Focal. LKO is measured in person-years. Japanese rates of 1947 and 2019.

**Figure A-1:**
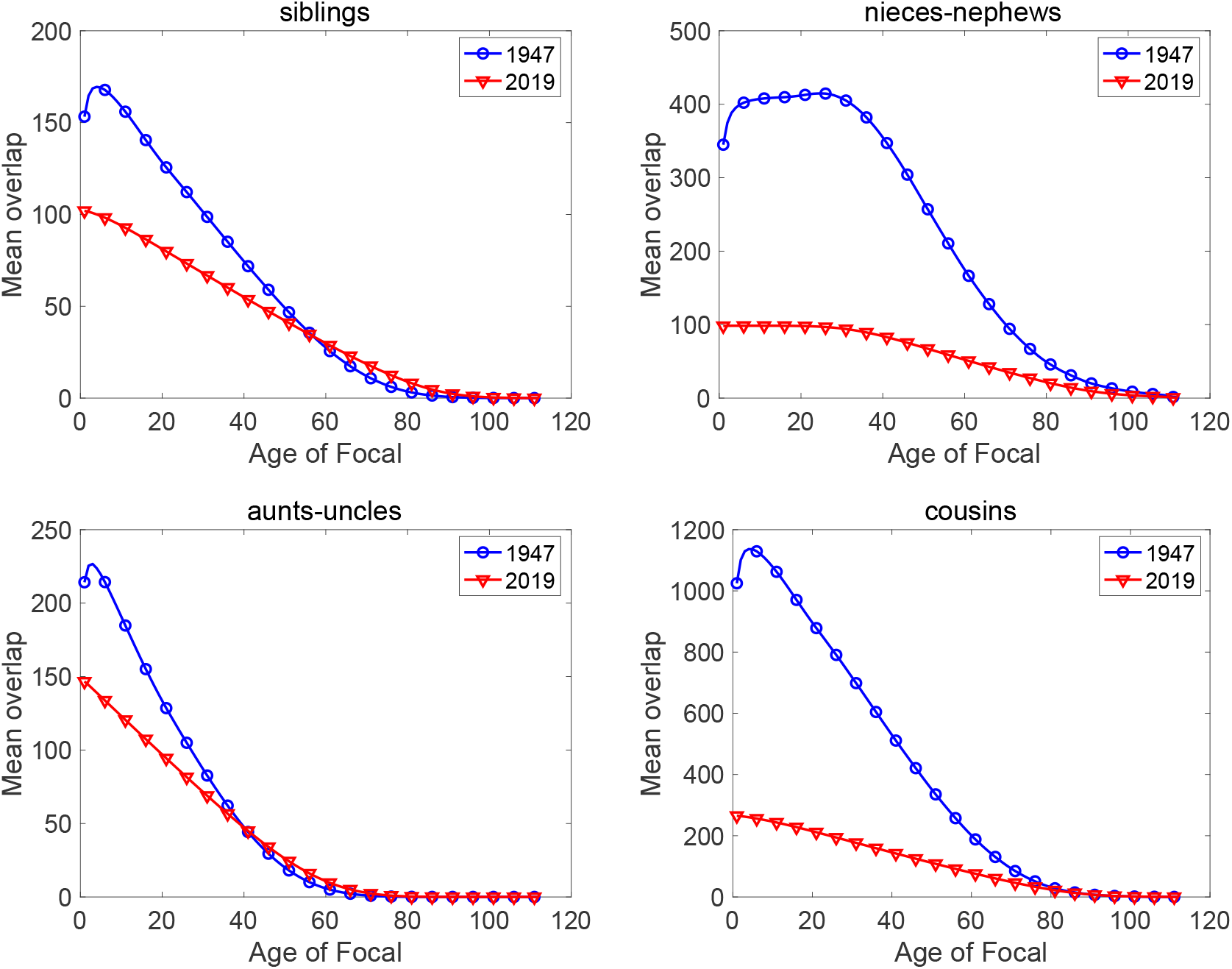
(**Part 2**) Mean remaining LKO with each type of kin, both sexes combined, as a function of the age of Focal. LKO is measured in person-years.Japanese rates of 1947 and 2019.

### A.2 Overlap with female and male kin

**Figure A-2:**
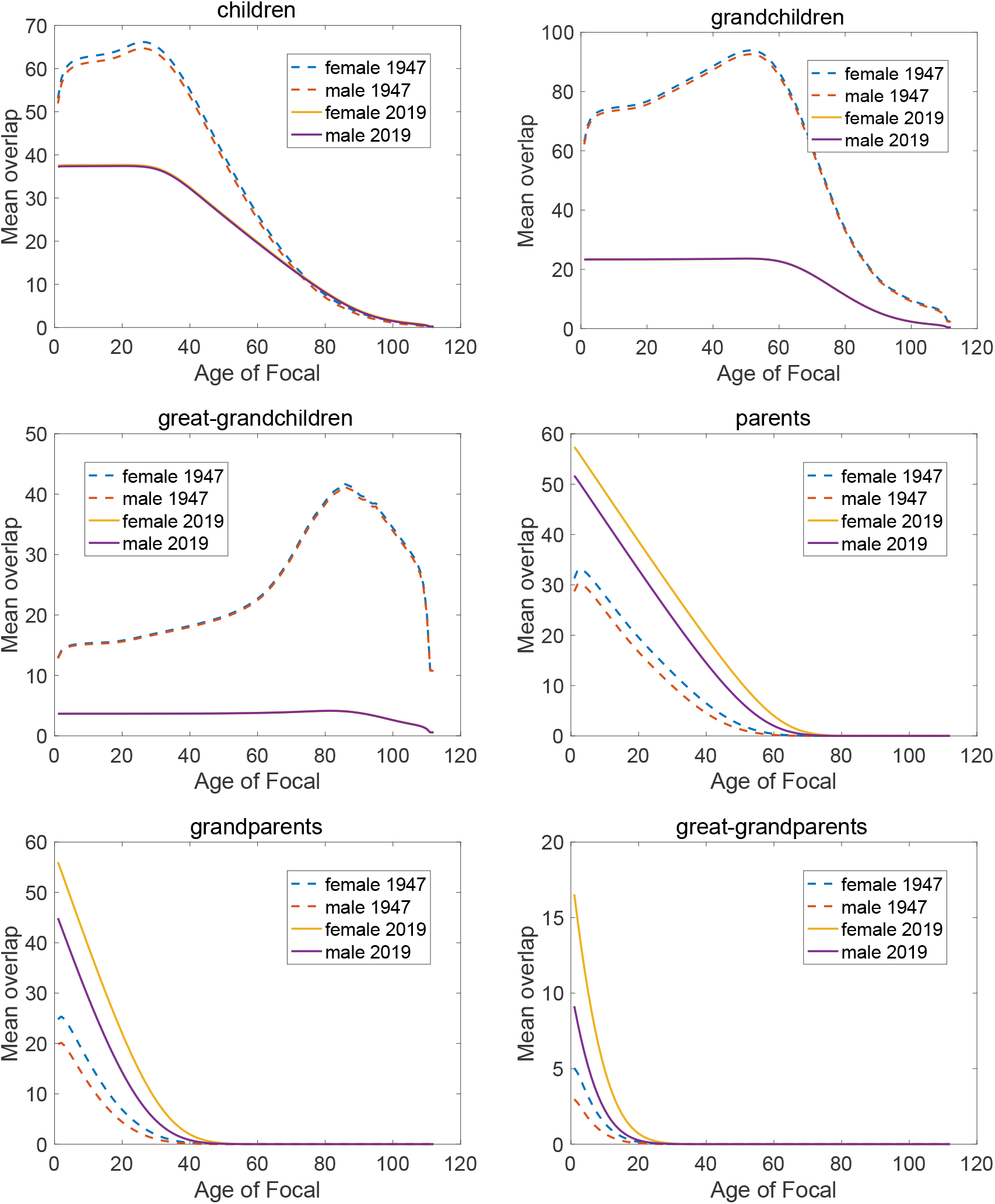
(**Part 1**) Mean remaining LKO with male and female members of each type of kin, as a function of the age of Focal. LKO is measured in person-years. Japan rates for 1947 and 2019.

**Figure A-2:**
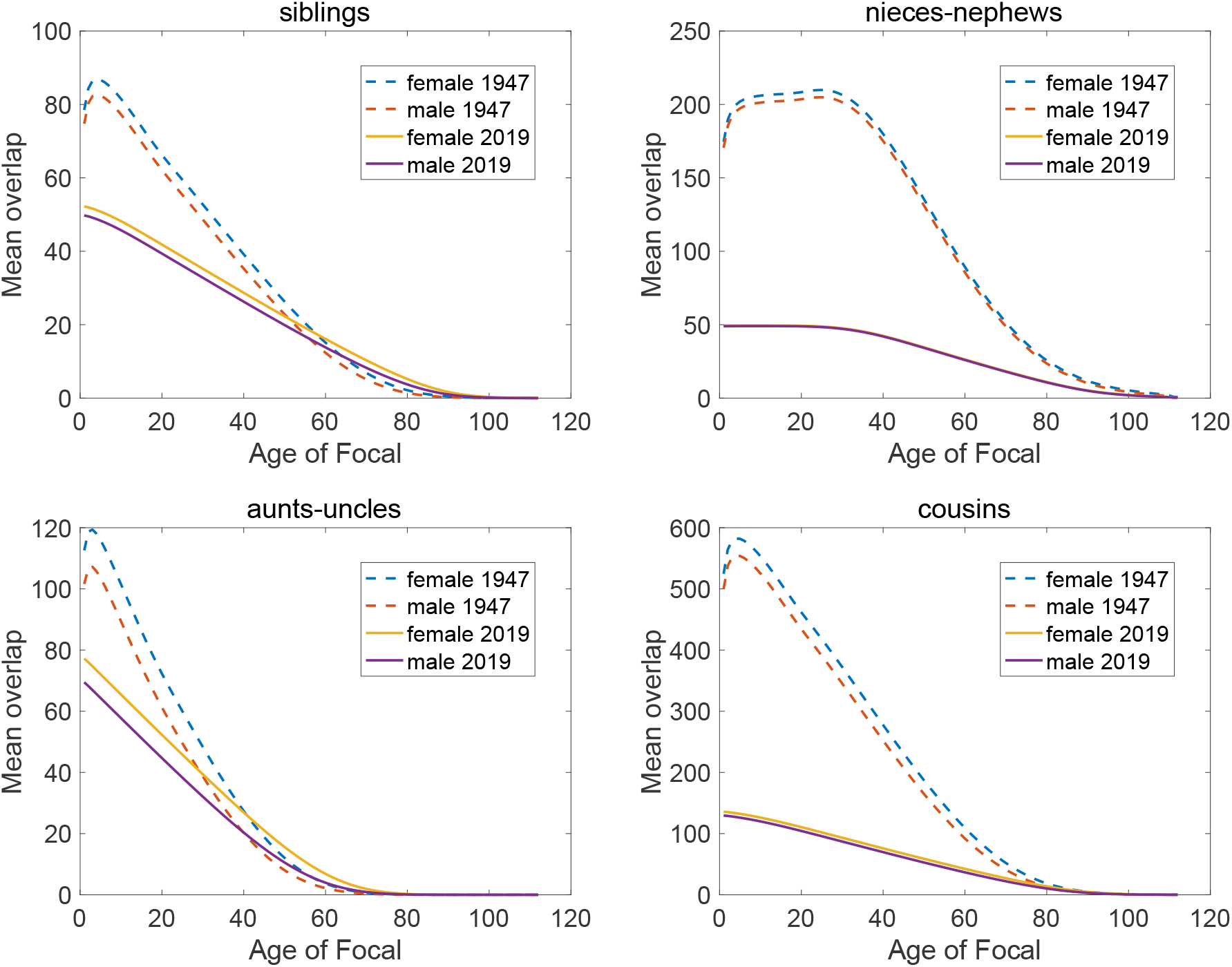
(**Part 2**) Mean remaining LKO with male and female members of each type of kin, as a function of the age of Focal. LKO is measured in person-years. Japan rates for 1947 and 2019.

### A.3 Overlap with at least one kin

**Figure A-3:**
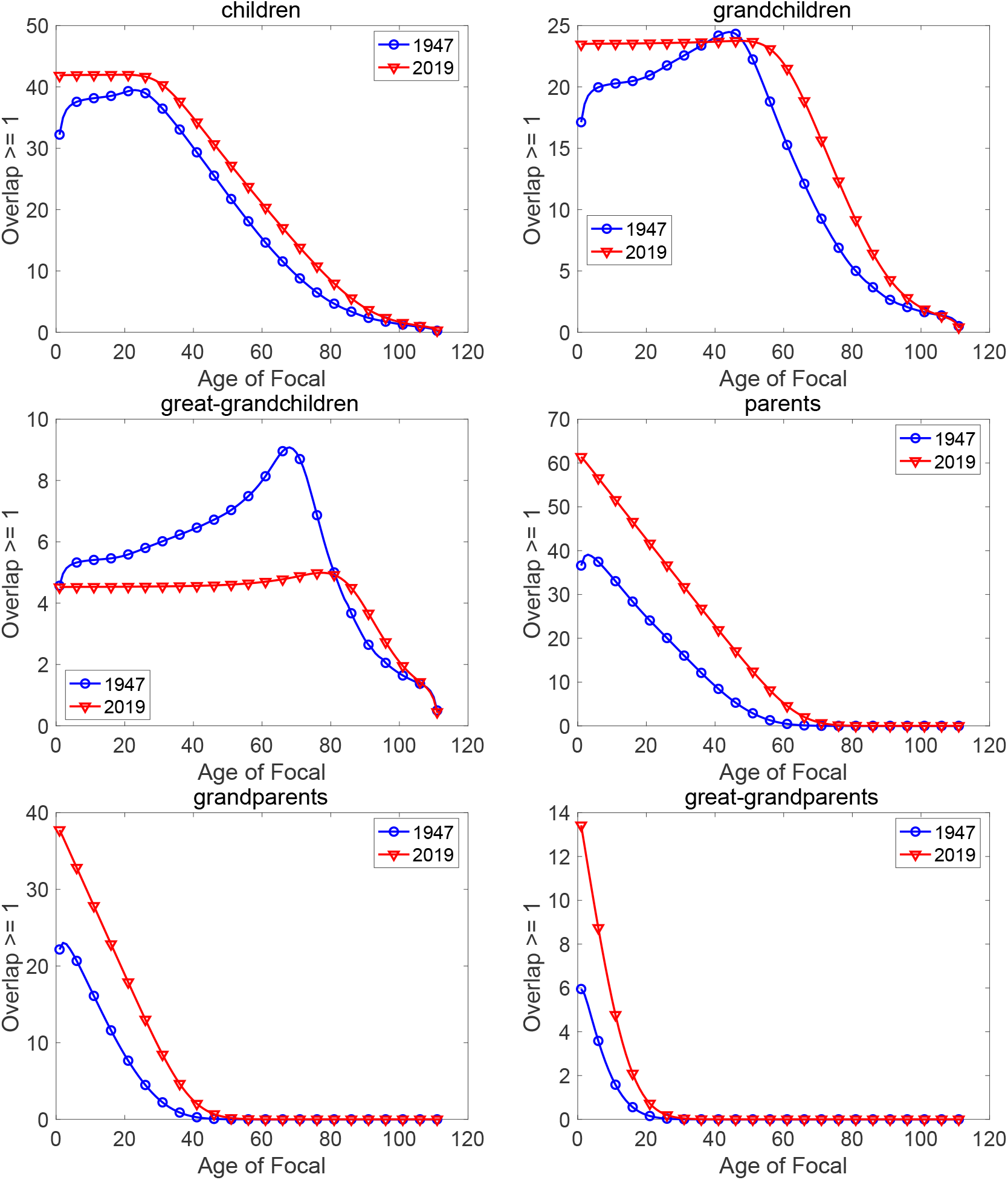
(**Part 1**.) Mean remaining LKO with at least one individual of each type of kin, as a function of the age of Focal. LKO is measured in years. Both sexes combined. Japan rates for 1947 and 2019.

**Figure A-3:**
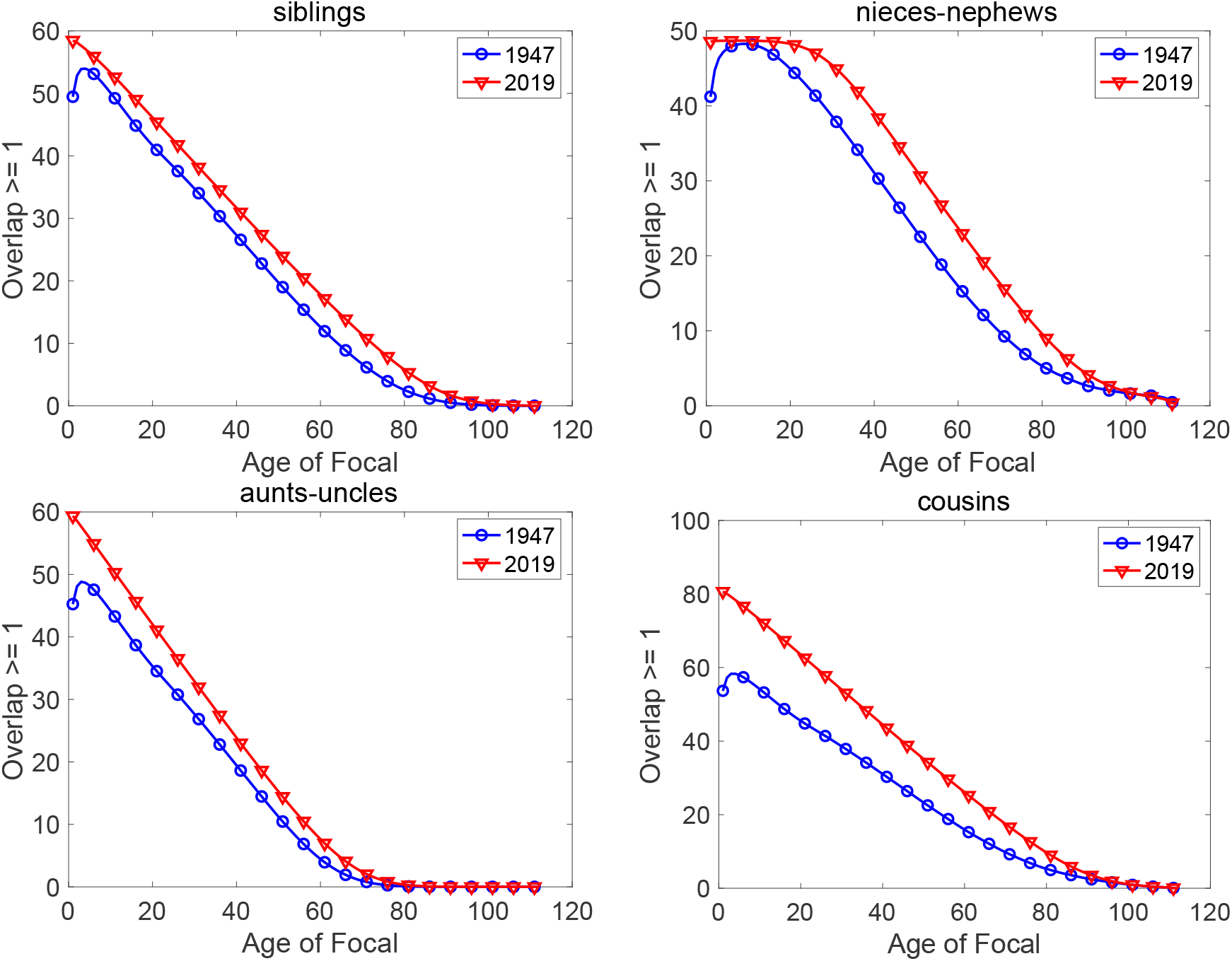
(**Part 2**) Mean remaining LKO with at least one individual of each type of kin, as a function of the age of Focal. Both sexes combined. LKO is measured in years. Japan rates for 1947 and 2019.

### A.4 Standard deviation of kin overlap

**Figure A-4:**
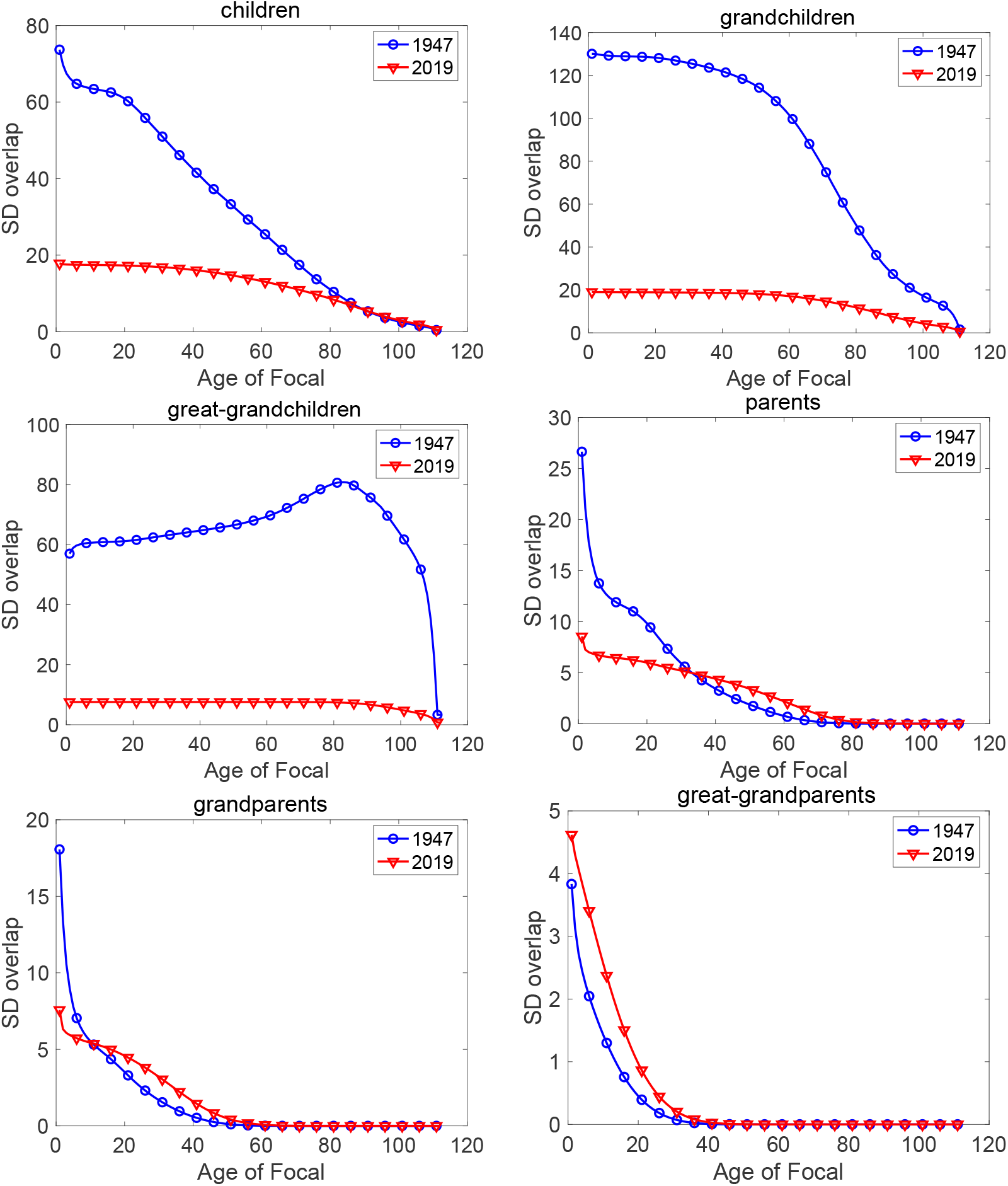
(**Part 1**) Standard deviation (SD) of remaining LKO with each type of kin, both sexes combined, as a function of the age of Focal. The SD of LKO is measured in person-years. Japan rates for 1947 and 2019.

**Figure A-4:**
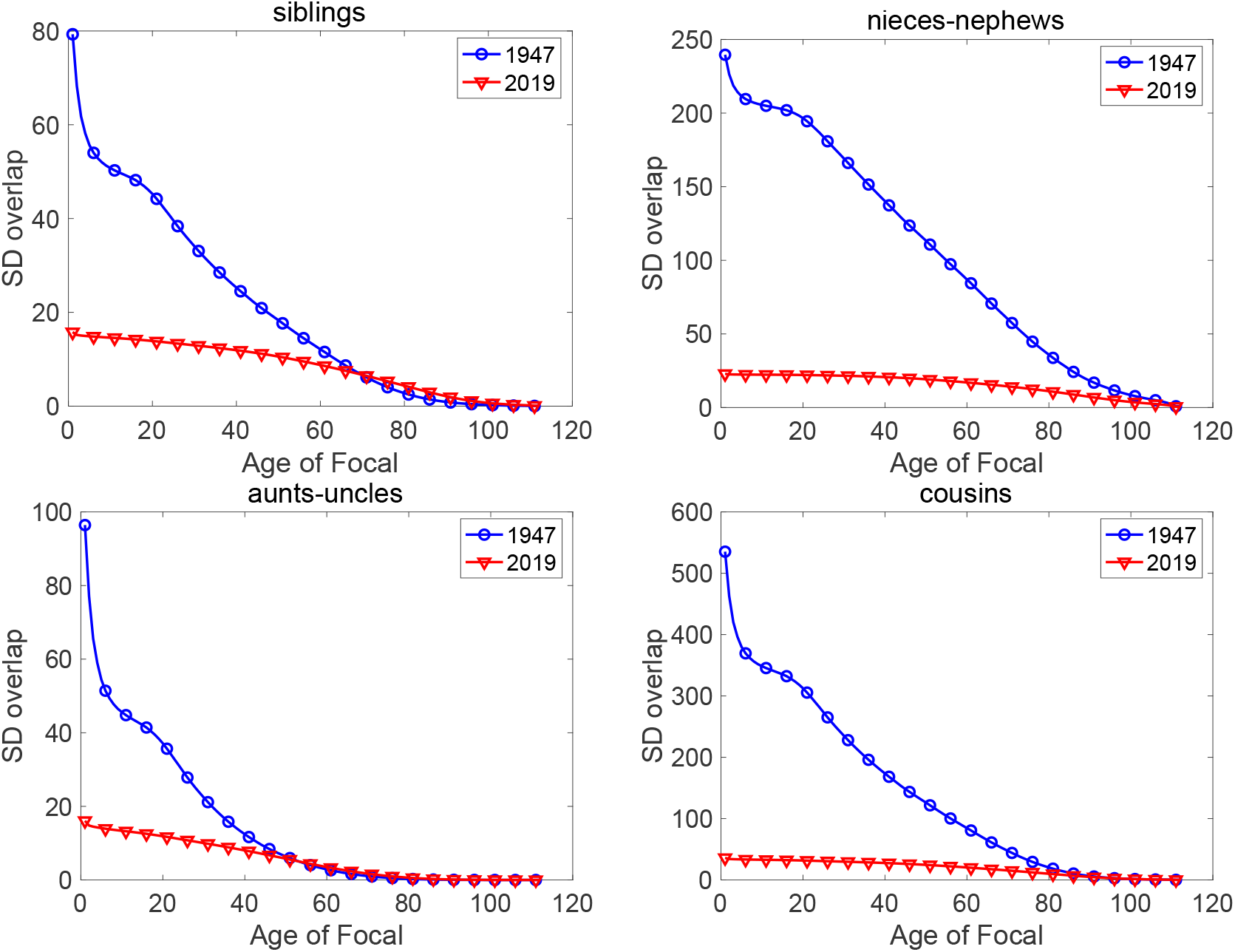
(**Part 2**) Standard deviation (SD) of remaining LKO with each type of kin, both sexes combined, as a function of the age of Focal. The SD of LKO is measured in person-years.Japan rates for 1947 and 2019.

### A.5 Prediction intervals for kin overlap

**Figure A-5:**
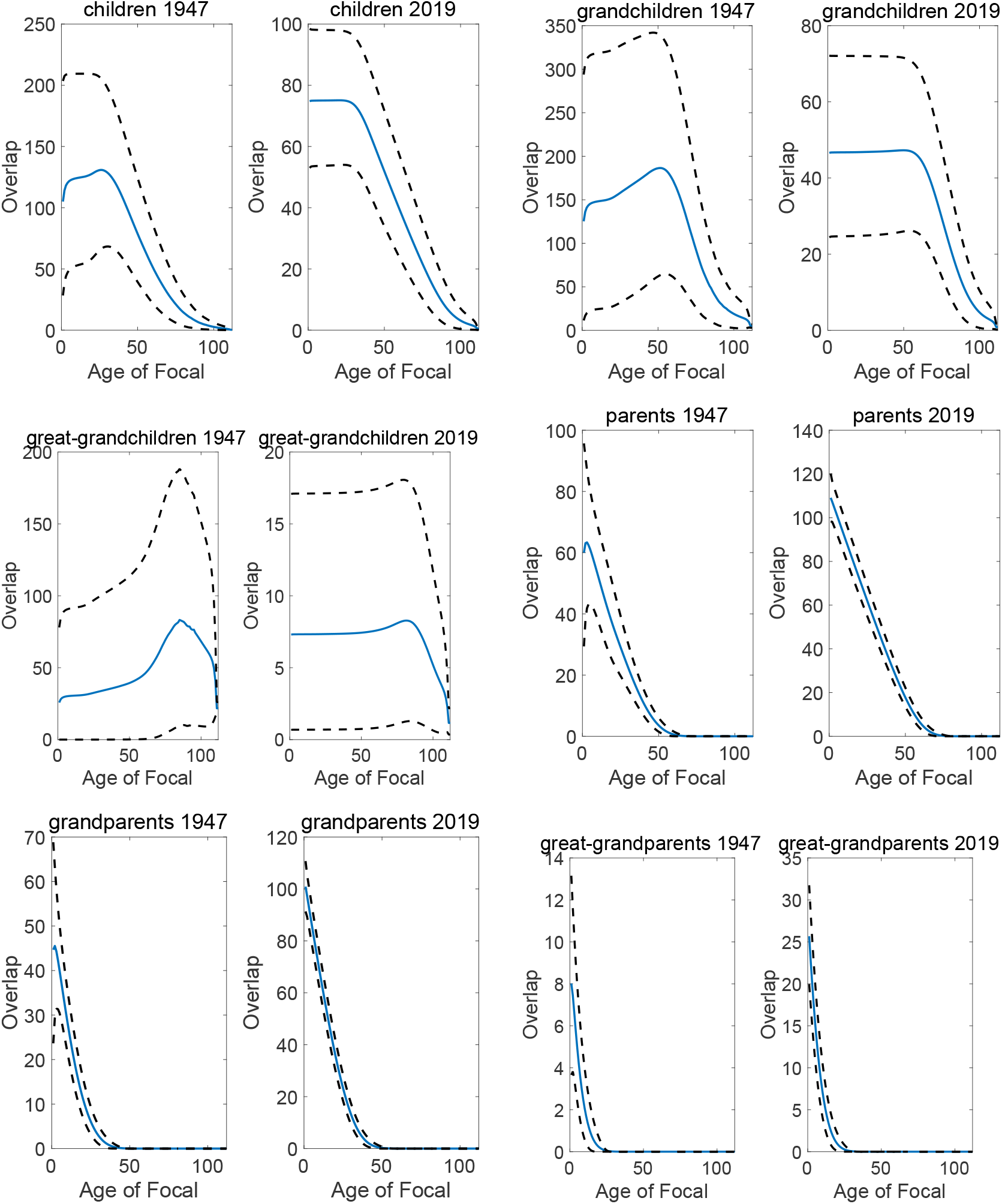
(**Part 1**) Expected remaining LKO with each type of kin, both sexes combined, with 90% prediction intervals calculated from a gamma distribution. LKO is measured in person-years. Japan rates of 1947 and 2019.

**Figure A-5:**
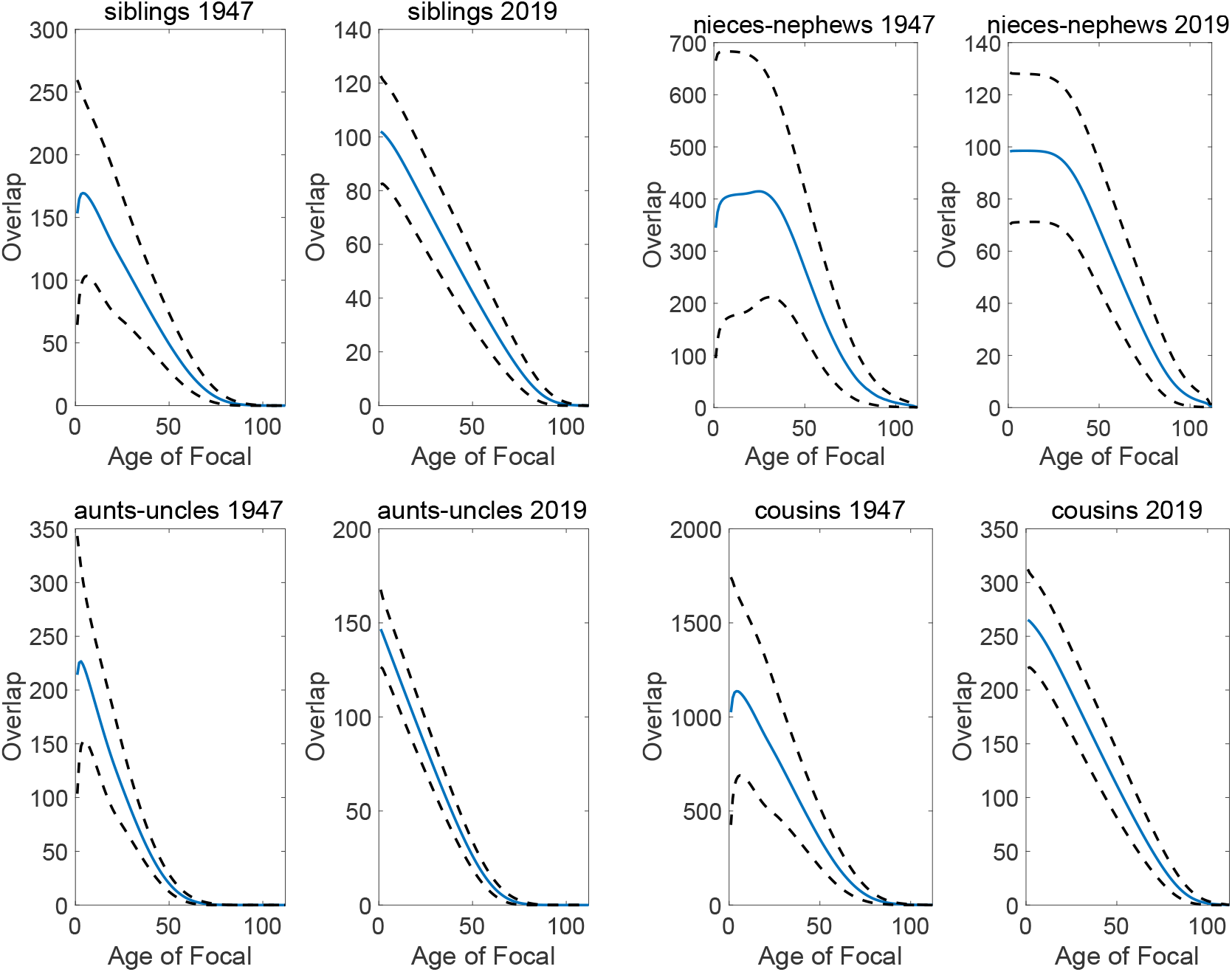
(**Part 2**) Expected remaining LKO with each type of kin, both sexes combined, with 90% prediction intervals calculated from a gamma distribution. LKO is measured in person-years. Japan rates of 1947 and 2019.

### A.6 Retrospective mean lifetime overlap with kin

**Figure A-6:**
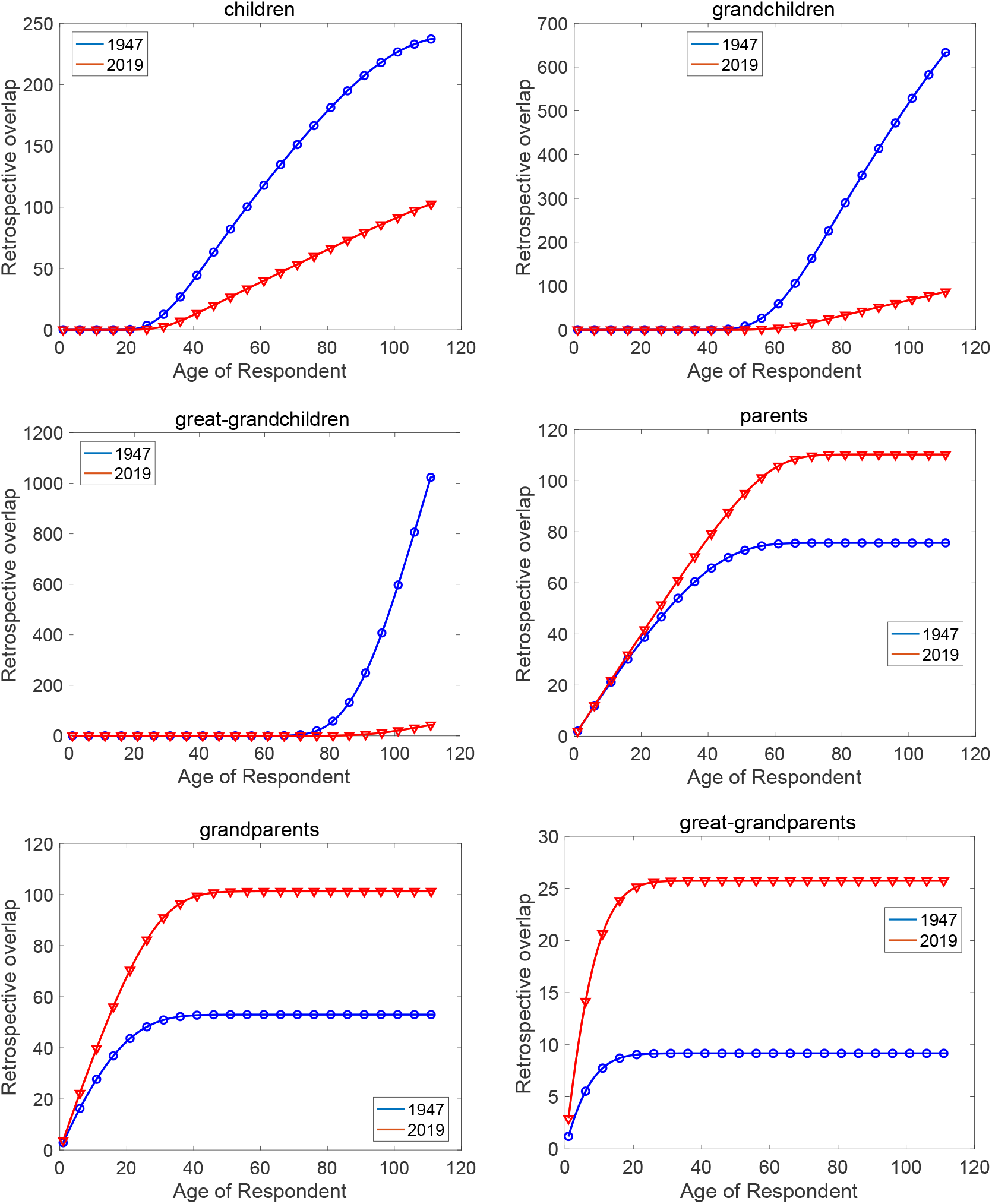
(**Part 1**) Mean retrospective LKO with each type of kin, both sexes combined, as a function of the age of Respondent. LKO is measured in person-years. Japanese rates of 1947 and 2019.

**Figure A-6:**
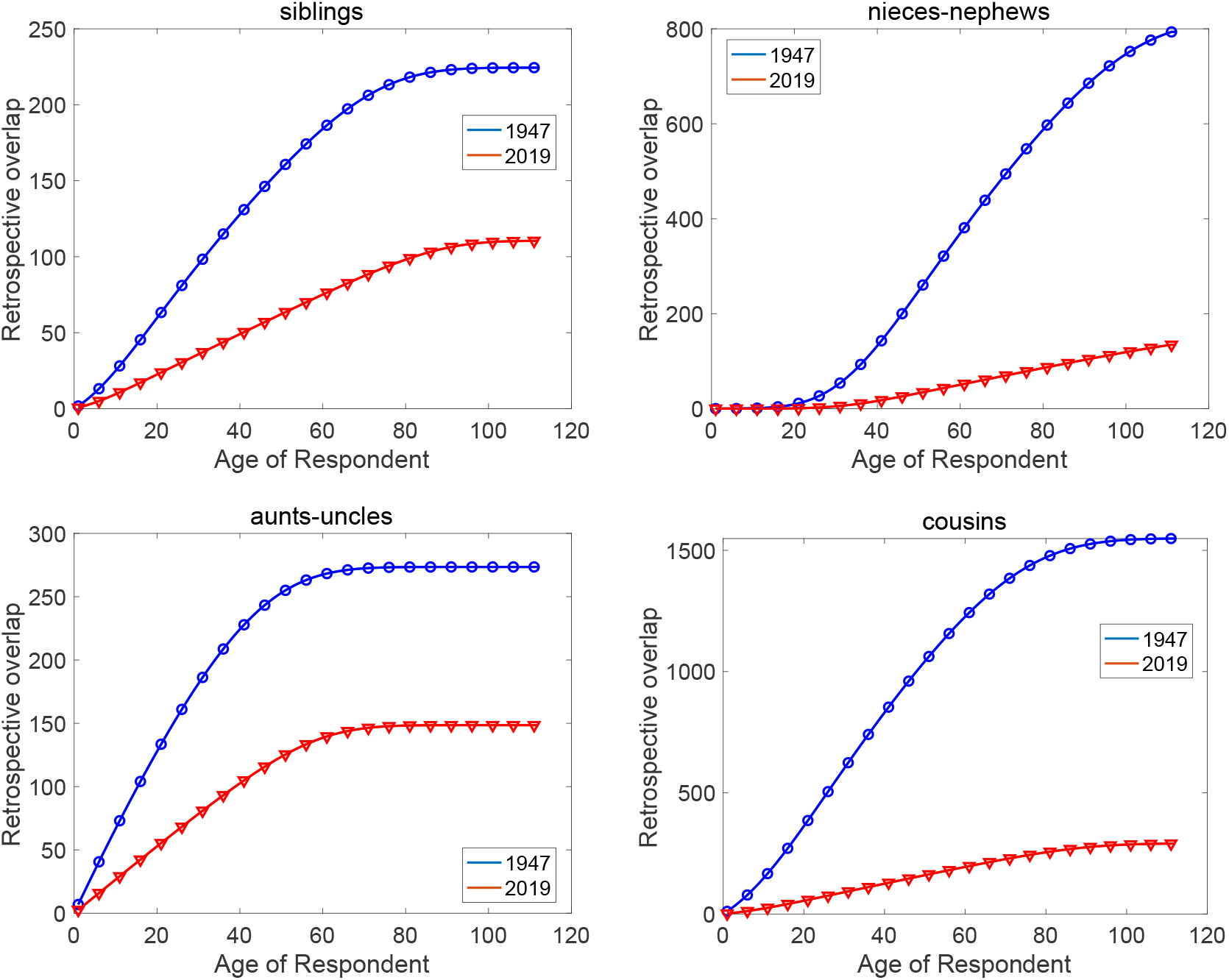
(**Part 2**) Mean retrospective LKO with each type of kin, both sexes combined, as a function of the age of Respondent. LKO is measured in person-years. Japanese rates of 1947 and 2019.

## Appendix B Projected mean kin numbers: Japan 1947 and 2019

**Figure B-1:**
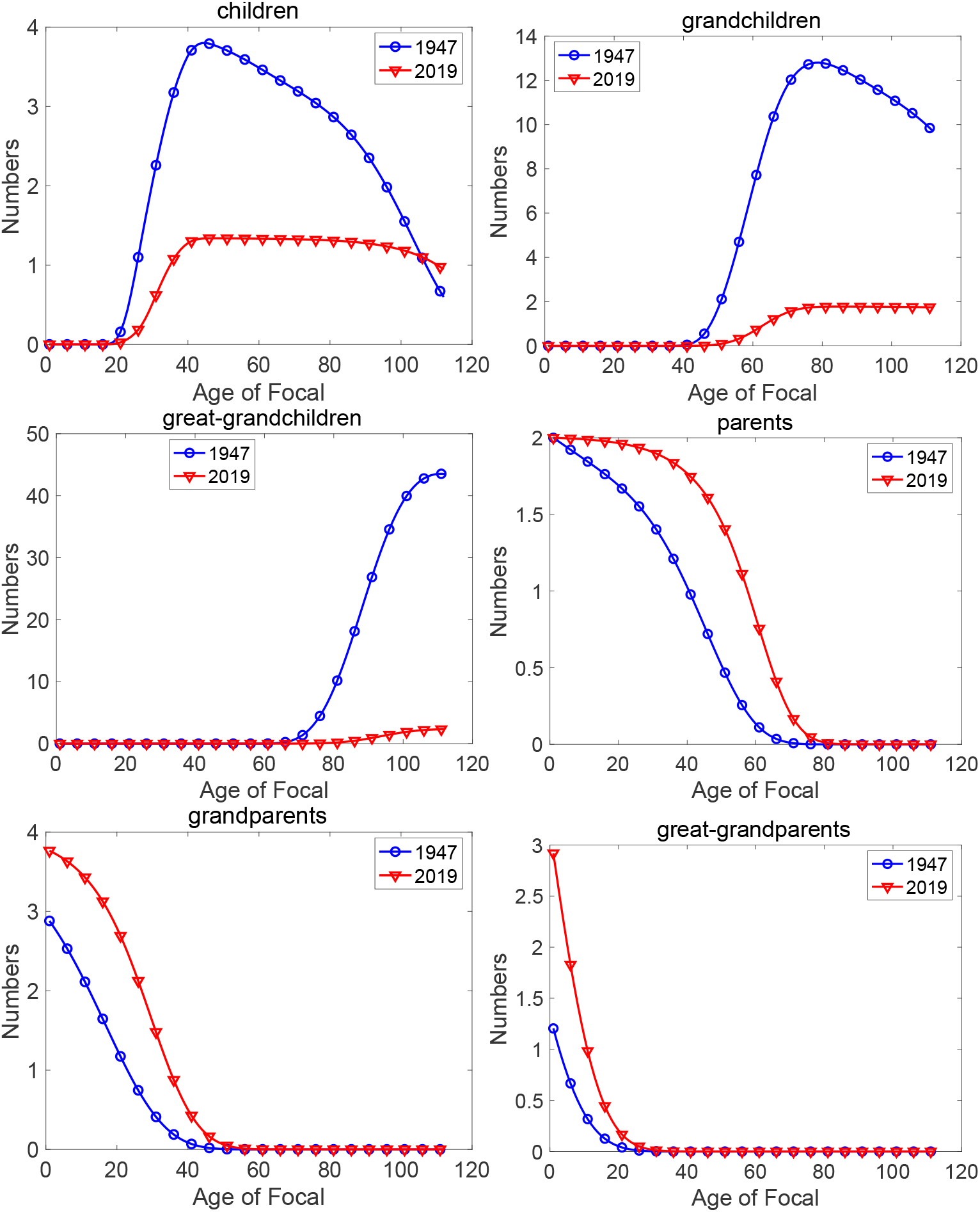
(**Part 1.)** Expected numbers of each type of kin as a function of the age of Focal, for Japan under 1947 and 2019 rates.

**Figure B-1:**
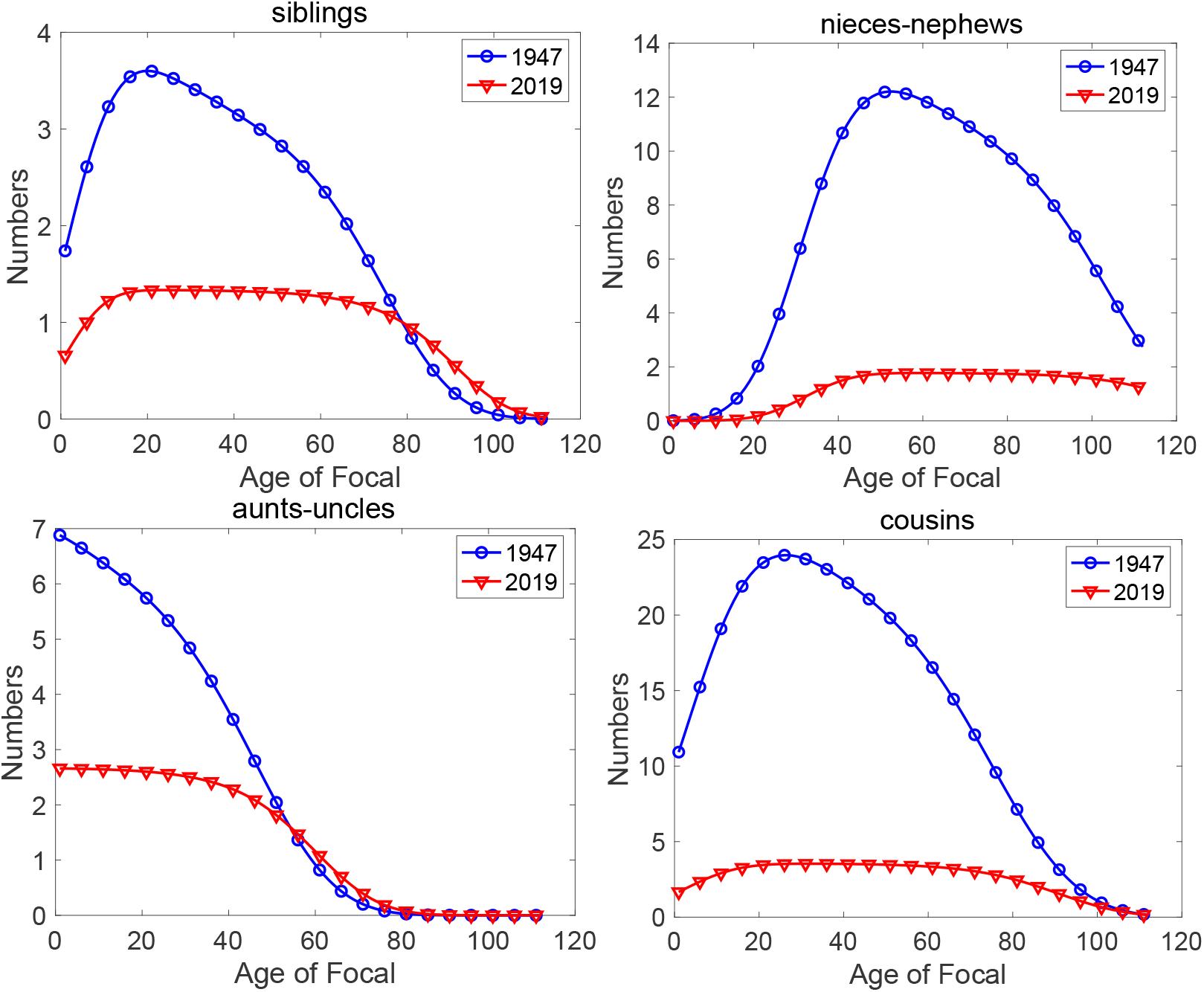
(**Part 2.)** Expected numbers of each type of kin as a function of the age of Focal, for Japan under 1947 and 2019 rates.

1 This notation is an intentional analogy to terms such as lifetime reproductive success (LRS), lifetime reproductive output (LRO), net reproductive rate (NRR), total fertility rate (TFR), and healthy life expectancy (HE or HALE). The calculation of these indices all begin with an age- or stage-specific property (fertility, healthy life, and so on) and then integrate that property over all or part of a lifetime.

2 If neither sex-specific mortality or fertility schedules are available, female rates can be applied to both sexes(the androgynous approximation) to obtain male and female kin numbers (Caswell, 2022).

3 Markov chain models can also describe mutistate demography in which the states correspond to stages (e.g., health conditions or parity classes) in combination with age classes. We do not consider these further here, but it is important that the methods we present apply to them as well.

4 The transition matrix is written in column-to-row orientation, to agree with the orientation of population projection matrices.

5 This is the first use of MCWR methods for kinship analysis. This analysis of Markov chains was introduced by Howard (1960) as the basis for stochastic dynamic programming (e.g., Puterman, 1994; Sheskin, 2010). Caswell (2011) extended the theory to include random rewards and demographic models; since then MCWR models have been applied to lifetime reproductive output (Caswell, 2011; van Daalen and Caswell, 2015; van Daalen and Caswell, 2017), evolutionary biodemography (van Daalen and Caswell, 2024), income and expenditures (Caswell and Kluge, 2015), and healthy longevity (Caswell and Zarulli, 2018; Owoeye, Oseni, and Gayawan, 2020; Caswell and van Daalen, 2021; Zarulli and Caswell, 2024).

6 This formulation assumes that the overlap experienced at each age is identical regardless of the transition made, except for the transition to death.

7 Perhaps a more general analysis would define a utility function for the number of, say, cousins. The age-specific overlap with all cousins is a utility function with constant returns; each additional cousin increases the overlap in that year by 1.The overlap with at least one cousin is an extreme case of a utility function with diminishing returns; once a single cousin is present, additional cousins add nothing to the overlap in that year. A utility function could easily be devised that would exhibit either increasing or decreasing returns. This is an open research problem.

8 Please do not become confused between the Poisson or Binomial distribution of kin numbers and the Bernoulli distribution of the presence of at least one kin.

9 This intervals are not confidence intervals surrounding an estimated mean, but rather intervals capturing a specified percentage of the distribution of LKO. The mean itself is calculated exactly from the mortality and fertility schedules and is not subject to sampling variation.

10 The product appears in various contexts as a measure of simultaneous occurrence. It corresponds to the AND operator in symbolic logic (two statements being simultaneously true) and to the intersection operator in set theory (for elements to belong simultaneously to two sets). In the algebra of events underlying probability theory, the product of two events describes the occurrence of both events (Ré nyi, 1970).

